# Chemically-informed coarse-graining of electrostatic forces in charge-rich biomolecular condensates

**DOI:** 10.1101/2024.07.26.605370

**Authors:** Andrés R. Tejedor, Anne Aguirre Gonzalez, M. Julia Maristany, Pin Yu Chew, Kieran Russell, Jorge Ramirez, Jorge R. Espinosa, Rosana Collepardo-Guevara

## Abstract

Biomolecular condensates composed of highly charged biomolecules like DNA, RNA, chromatin, and nucleic-acid binding proteins are ubiquitous in the cell nucleus. The biophysical properties of these charge-rich condensates are largely regulated by electrostatic interactions. Residue-resolution coarse-grained models that describe solvent and ions implicitly are widely used to gain mechanistic insights into the biophysical properties of condensates, offering transferability, computational efficiency, and accurate predictions for many systems. However, their predictive accuracy diminishes for charge-rich condensates due to the implicit treatment of solvent and ions. Here, we present the Mpipi-Recharged model, a residue-resolution coarse-grained model that improves the description of charge effects in biomolecular condensates containing disordered proteins, multi-domain proteins, and/or disordered RNAs. Mpipi-Recharged maintains the computational efficiency of its predecessor—the Mpipi model—by still treating solvent and ions implicitly, but improves its accuracy by incorporating a pair-specific asymmetric electrostatic potential informed by atomistic simulations in explicit solvent and ions. We show that such asymmetric coarse-graining of electrostatic forces is needed to recapitulate the stronger mean-field impact of associative interactions between opposite-charge pairs over the repulsion among equally charged pairs revealed by our atomistic simulations. Mpipi-Recharged shows excellent agreement with the experimental phase behavior of highly charged systems, capturing subtle effects challenging to model without explicit solvation, such as the impact of charge blockiness, stoichiometry changes, and salt concentration variation. By offering improved predictions for charge-rich biomolecular condensates, Mpipi-Recharged extends the computational tools available to investigate the physicochemical mechanisms regulating biomolecular condensates.

## I. INTRODUCTION

Biomolecular condensates are ubiquitous intracellular assemblies formed via phase separation of biomolecular mixtures^1–4^. Multivalency is a universal feature of phase-separating biomolecules, as it enables them to form dynamic, percolated networks that ensure the thermodynamic stability of the condensate^5^. The greater the biomolecular valency, the denser the network of inter-molecular connections that biomolecules can form, and the greater the enthalpic gain for condensate formation^5^. Despite valency being universal to phase-separating biomolecules, the dominant types of intermolecular interactions that different multivalent biomolecules establish to decrease the free energy of their condensates are highly varied. These interactions depend not only on the chemical makeup and structural properties of the biomolecules themselves but also on those of their binding partners and on the parameters of the solution, such as salt concentration^6–8^, temperature^9,10^ and pH^11^. Moreover, the presence of molecular crowding agents^12,13^, changes in post-translational modifications (PTMs)^14–16^, and fluctuations in metabolite concentrations^17–19^ are also potent modulators of the inter-molecular interactions inside condensates. This capacity for regulation highlights the ability of condensates to exhibit specificity and selectivity in their composition and biophysical properties.

In some condensates, the essential multivalent biomolecules are rich in aromatic residues and arginines, such as the RNA-binding proteins with prion-like domains found in cytoplasmic stress granules^2,20–23^. As a result, in such systems, π–π^9,24–28^ and cation– π^6,24,27,29–31^ interactions are the chief drivers of biomolecular phase separation, at least at physiological salt concentrations^9,24,25,27,32,33^.

In contrast, in condensates filled mostly with charge-rich molecules, charge–charge interactions can be the dominant regulators. This is observed, for instance, in complex coacervates^7,34–36^ and in most nuclear condensates^37–39^. These condensates may contain mixtures of chromatin^40,41^, nucleosomes^37,42^, chromatin–binding proteins^43^, negatively charged polyelectrolytes— e.g., DNA^37–39,44^, RNA^45–51^, and IDRs or IDPs rich in glutamic and/or aspartic acids (e.g., prothymosin *α*)^52^— and positively charged polyelectrolytes—e.g., IDRs or IDPs rich in arginine and/or lysine (e.g., the histone tails of nucleosomes and the H1 protein)^7,34–36,52–55^. Additionally, they may contain polyampholytes, which are IDRs or IDPs like the N-tail of DDX4, possessing well-balanced distributions of both positively and negatively charged residues, and a low net charge per molecule^56^.

Understanding the molecular and biophysical mechanisms by which charge–charge interactions impact the properties of condensates is highly desirable but significantly challenging. Some charge-rich proteins, such as *α*-synuclein, undergo phase separation only in the presence of molecular crowders or at high salt concentration (where repulsive electrostatic interactions are sufficiently screened)^57–59^. Other charge-rich biomolecules, phase separate only as part of multi-component systems by establishing associative heterotypic interactions commonly of electrostatic nature (e.g. the mixture of H1 and prothymosin *α*)^7,45^.

Computer simulations represent a powerful tool to probe the molecular mechanisms by which charge-rich biomolecules regulate the physicochemical properties of their biomolecular condensates. Among these tools, transferable residue-resolution coarse-grained models for biomolecular phase separation^24,29,60–70^, such as the Mpipi family^24,64^, the CALVADOS family^61,62^, the HPS family^29,60,66,71^, and MOFF^72^ have gained notoriety as they can effectively balance computational efficiency with adequate physicochemical accuracy. These models represent amino acids with a single bead, and define residue-pair interactions using a combination of short-range potentials—to model non-electrostatic interactions and excluded volume (e.g. the Ashbaugh-Hatch version^73^ of the 12-6 Lennard-Jones potential^74^ for HPS and CALVADOS, and the Wang– Frenkel potential^75^ for Mpipi^24^)—and long-range Debye– Hückel potentials—to represent salt-screened electrostatic interactions. Emergent properties of biomolecular solutions–such as temperature-vs-density phase diagrams^6,29,62,76,77^, saturation concentrations^76,78,79^, relative condensate stability^24,61,78,80^, viscoelastic properties^50,78,81–83^, and multiphase organization–are estimated using such models via Direct Coexistence Molecular Dynamics simulations^84^ or constant-pressure simulations of the condensed phase^85,86^. Notably, simulations of single-molecule IDRs and of protein solutions using the CALVADOS^61–63^ and the Mpipi models^24,64^ have achieved excellent agreement with experiments in measuring IDR ensemble properties, phase diagrams, and saturation concentrations^24,27,64,77,87^.

Some innovative aspects of the Mpipi coarse-grained model^24^ include: (a) the use of the Wang–Frenkel potential^75^ to describe non-electrostatic associative interactions, (b) a parameterization derived from a combination of atomistic simulations of amino acid pairs^24^ and bioinformatics data^88^, and (c) the replacement of the commonly used Lorentz–Berthelot combination rules— used to define the strength of non-electrostatic interactions among amino acid pairs from the hydrophobicity scales of the pure amino acids^29,60–62,71^—for amino-acid-pair-specific parameters. These three features combined enhance the computational efficiency of the model and the parameterization flexibility. Additionally, they allow the model to accurately capture the stronger role of π-based contacts compared to non-π interactions, the significantly stronger associative contacts established by arginines over lysines, and the significantly stronger bonds that arginine establishes with aromatic residues (cation-π) over non-aromatic species. As a result, the predictions of the Mpipi model for the temperature-vs-density phase diagrams of the variants of the IDR of the hnRNPA1 protein^24^ are in near-quantitative agreement with experiments^9^. Furthermore, the predictions of Mpipi regarding the relative change in thermodynamic stability as a function of amino-acid mutations for several prion-like domain proteins are consistent with qualitative experimental trends^27^. More recently, the Mpipi-GG model^64^ refined some of the Mpipi Wang–Frenkel parameters, including stronger glycine–glycine and glycine– serine interactions, weaker aromatic–charge interactions, and generally stronger interactions for alanine, leucine, and isoleucine. Subsequently, the deep learning algorithm ALBATROSS, developed using Mpipi-GG, was used to predict ensemble dimensions of 137 IDRs in excellent agreement with experiments^64^.

Regarding the description of charge–charge interactions, even the most successful residue-resolution coarse-grained models, such as Mpipi^24,64^ and CALVADOS^61,62^, have certain limitations^6,7,37,66,68–70,89,90^. These limitations arise from the challenge of accurately capturing the impact of water and ions in the regulation of electrostatic forces in condensates, while describing these components implicitly in favour of computational efficiency. For example, the release of ions and water upon condensation is expected to play a crucial and complex role in regulating processes such as the salt-dependent modulation of protein condensate stability ^6,7^, the RNA-driven reentrant phase behavior of protein condensates^47^, the impact of charged patterning on condensates stability and preferential partitioning^91–93^, and complex coacervation^45^.

In this work, we present the Mpipi-Recharged model, which improves the description of charge–charge interactions in biomolecular phase behaviour, while maintaining the accuracy and performance of its predecessor^24,27^.. The new Mpipi-Recharged model describes charge–charge interactions with an assymmetric pairspecific Yukawa potential designed to recapitulate the results from our atomistic potential of mean force simulations in explicit solvent and ions. These atomistic simulations reveal that, at the mean field level, the relative strength of electrostatic association and repulsion among charged amino acid pairs is substantially different; i.e., after averaging out over all degrees of freedom except for the inter-molecular separation, the associative interactions between opposite-charge pairs are significantly stronger than the repulsive ones among equally-charge pairs. We show that our pair-specific Yukawa potential successfully captures such asymmetry and allows Mpipi-Recharged to reproduce the phase behaviour of highly charged proteins and complex coacervates.

We thoroughly validate the Mpipi-Recharged model against a range of experimental measurements, including single-molecule radius of gyration, critical solution temperatures, and saturation concentrations. Additionally, we show that the Mpipi-Recharged model accurately captures experimental trends of various difficult-to-model highly charged systems, including salt-dependent phase diagrams, the impact of charge blockiness on the relative stability of condensates, and the stoichiometry thresholds observed for the various regimes of the RNA-driven reentrant phase behaviour for various RNA-binding proteins. Overall, the Mpipi-Recharged model exemplifies how incorporating a pair-specific asymmetric potential for electrostatic forces significantly improves the description of the phase behaviour of highly charged systems without the need for explicit solvent and ions.

## II. RESULTS AND DISCUSSION

### A. Atomistic simulations suggest asymmetric pair-specific coarse-graining of electrostatic forces

We first quantify the relative strengths of interactions between different pairs of charged amino acids by conducting atomistic umbrella sampling Molecular Dynamics simulations. These simulations were performed with fixed amino acid orientations in explicit solvent and ions at 298 K, employing the a99sb-disp/JC-SPC/E-ion/TIP4P/2005 force field combination^94,95^ (see Methods for details). From the umbrella sampling simulations, we calculate the potential of mean force of the amino pairs as a function of their pairwise centre-of-mass distance^6,18,28,77,79,96–100^. To elucidate the differences in attractive vs. repulsive forces, we computed potential of mean force calculations for all the ten unique combinations of the four amino acids with total charges (*q*) equal to +1e or -1e at pH=7. These amino acids are arginine (R, *q* = +1e), lysine (K, *q* = +1e), glutamic acid (E, *q* = −1e), and aspartic acid (D, *q* = −1e). The combinations analyzed are aspartic acid–aspartic acid (D–D), glutamic acid–aspartic acid (E–D), glutamic acid–glutamic acid (E–E), lysine–lysine (K–K), lysine–arginine (K–R), arginine–arginine (R–R), arginine–glutamic acid (R–E), lysine–glutamic acid (K–E), lysine–aspartic acid (K–D), and arginine–aspartic acid (R–D) (Fig. 1). Each of these potential of mean force calculations describe the change in free energy that the amino acids in a given charged pair experience as they approach one another from infinity. This energy has been averaged out over all degrees of freedom of the system except for the one-dimensional intermolecular pairwise separation, which we use as the reaction coordinate. The minimum value of these potential of mean force calculations will be referred to herein as the ‘binding free energy’.

**FIG. 1.**
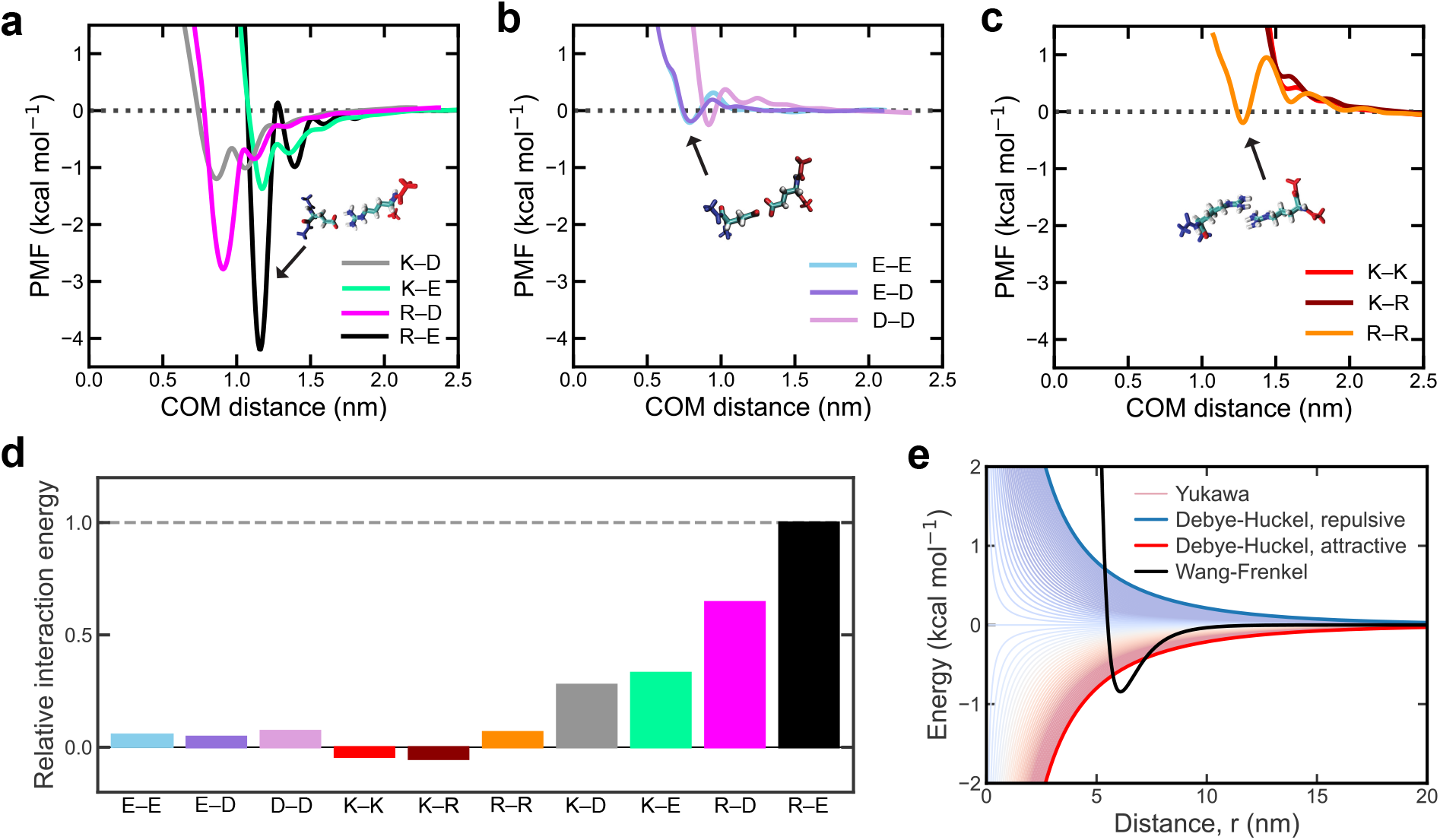
Free-energies of binding between ±1 charged amino acid pairs reveal a stronger role of associative versus repulsive electrostatic forces. Potential of Mean Force (PMF) computed from umbrella sampling atomistic Molecular Dynamics simulations of charged amino acid pairs. The center-of-mass (COM) distance between the amino-acid pair was used as the reaction coordinate. The simulations were performed with the a99SB-disp all-atom force field^94^ at T=298 K and 150 mM NaCl salt concentration. The potential of mean force calculations shown are for: **a**. Positive–negative (Black: glutamic acid–arginine, E–R, Green: glutamic acid–lysine, E–K, Grey: lysine–aspartic acid, K–D, and Magenta: arginine–aspartic acid, R–D), pairs of charged residues. **b**. Negative–negative (pink: aspartic acid–aspartic acid, D–D, Purple: glutamic acid–aspartic acid, E–D, and blue: glutamic acid–glutamic acid, E–E), and **c**. Positive-positive (Red: lysine–lysine, K–K, Maroon: lysine–arginine, K–R, and Orange: arginine–arginine, R–R) pairs of charged residues. **d**. Comparison of the free-energy of binding among charged amino acids relative to the R–E interaction. The values for the interaction are obtained from the minimum value of the corresponding potential of mean force curve in panels **a**-**c. e**. Schematic depiction of the potentials used in the Mpipi-Recharged model for hydrophobic/dispersive interactions (Wang-Frenkel potential; black thick curve), and electrostatic interactions (Yukawa potential; thin curves ranging from blue to red). Thick blue and red curves represent the standard profile of a Coulombic-like electrostatic screened potential (among species with charges equal to ±1) for repulsive and attractive interactions, respectively.

Our simulations reveal that, at small molecular separations, pairs of oppositely charged amino acids (i.e., arginine–glutamic acid, arginine–aspartic acid, lysine– glutamic acid and lysine–aspartic acid) experience a substantial reduction in their free energy due to the formation of associative electrostatic interactions (Fig. 1a). This result is consistent with our previous simulations^6,24^ and with the critical role of associative electrostatic interactions in promoting the stability of biomolecular condensates observed experimentally^6,7,45,46,101,102^. Additionally, our simulations show that, despite having the same charge, arginine exhibits a greater propensity than lysine to form associative interactions with negatively charged amino acids (Fig. 1a). The association of arginine with both glutamic acid and aspartic acid results in a notable free energy decrease—about −4 kcal/mol for arginine–glutamic acid and −3 kcal/mol for arginine– aspartic acid. In contrast, lysine shows a significantly smaller free energy decrease—up to −1.5 kcal/mol— upon binding to either glutamic acid or aspartic acid. The significantly more favourable binding free energies of arginine versus lysine with negative residues is consistent with experiments and simulations revealing a decreased phase separation propensity of protein solutions upon arginine to lysine mutation^24,27,103,104^ and the diverse properties of arginine-rich versus lysine-rich condensates^6,9,45,105^. These results also align well with the higher free energy of hydration for arginine compared to lysine^106^.

In stark contrast, the potential of mean force calculations for pairs of like-charged amino acids (negative– negative pairs in Fig. 1b and positive–positive pairs in Fig. 1c) indicate a significantly weaker interaction, when compared with those for positive–negative pairs. As the like-charged amino acids are brought closer to one another, the change in their free energy presents maxima of low height (less than +1 kcal/mol) signalling weak repulsion. For some of like-charged amino acid pairs, such arginine–arginine (Fig. 1c)) or the three negative– negative pairs (Fig. 1b), reducing the pairwise distance even more (to about 0.9-1.2 nm), reveals an additional weakly attractive interaction (a minimum of approximately −0.1 kcal/mol). We reasoned that such a minimal change in free-energy upon binding of like-charge amino acids, especially in comparison with the marked attraction observed among oppositely charged species, can be attributed to the following: (a) the partial reorientation of amino acid chains as the pairwise distance decreases, (b) alterations in ion solvation, and (c) changes in the translational entropy of ions and solvent. Specifically, as the pairwise centre-of-mass distance is reduced, the side-chains of opposite-charged amino acids, water molecules solvating them, and the ions around them reorganise themselves into configurations that maximize their Coulomb attraction, while in the cases of like-charged amino acids such reorganisation is aimed to minimize their repulsion.

### B. Mpipi-Recharged: a residue-resolution coarse-grained model with a pair-specific asymmetric electrostatic potential improves the description of charged-driven effects in condensates

Using atomistic models to study biomolecular phase separation in charged biomolecules would naturally account for the asymmetric contribution of electrostatic interactions between like-charged and oppositely-charged species we report in our potential of mean force calculations (Fig. 1); but, their high computational cost makes this impractical. Residue resolution coarse-grained models, like our model Mpipi^24^, offer a more efficient strategy to probe biomolecular condensation computationally. However, residue-resolution coarse-grained models for biomolecular phase separation ^24,29,60–66^ consider water and ions implicitly by invoking the Debye–Hückel mean-field approximation to describe electrostatic interactions^107^. The Debye–Hückel approximation is necessary to reduce the degrees of freedom of the system and make the simulations of condensates feasible; yet, it assumes that the effects of monovalent counterions in solution can be reduced to the screening of the mean electrostatic potential generated by the amino acid (or nucleotide) charges. Therefore, the approximation considers that the level of screening and, thus, the decay of charge–charge interactions with distance is the same in magnitude for amino acid pairs with identical absolute total charges (i.e., all pairs discussed in Fig. 1). That is because such decay is described by a Yukawa function that uses the same prefactor and screening length for all pairs of charged amino acids, regardless of their identity. Crucially, the notable differences in free-energy of binding among like-charged and opposite-charged amino acids pairs that we have observed atomistically (Fig. 1) cannot be recapitulated if the coarse-grained charge–charge interactions are described using the same Yukawa function parameters for all types of pairs. To overcome this limitation, here we developed a new residue-resolution coarse-grained model for biomolecular phase separation, named the Mpipi-Recharged model. The Mpipi-Recharged aims to improve the representation of charged biomolecules in our original Mpipi model while maintaining its exceptional performance for prion-like domain proteins^24,27^.

The Mpipi-Recharged model employs a Yukawa function with parameters that vary depending on the specific amino acid pair. Such a modification allows Mpipi-Recharged to consider that, at the coarse-grained level, attraction and repulsion should be described asymmetrically to compensate for the loss of explicit ions and water and for the aggressive mapping of the many charges that an amino acid carries atomiscally to just one charge centred on its alpha carbon when coarse-grained. To parameterize the pair-dependent Yukawa function, we use our atomistic potential of mean force calculations shown in Fig. 1 (see Methods for further details on the parameterization and model potentials). The resulting pair-specific parameters of the Yukawa functions are given in Table S3.

To evaluate the performance of the new Mpipi-Recharged model, we subject it to the same rigorous tests previously used for the original Mpipi model^24^. Specifically, we compare its performance against: (1) single-molecule radius of gyration (*R*_*g*_) experimental measurements of 46 different intrinsically disordered proteins (IDPs)^9,109–120^ from small-angle X-ray scattering (SAXS) and NMR (Fig. S3); and (2) experimental temperature-vs-concentration phase diagrams of hnRNPA1-LCD (A1-LCD) variants^9^ (sequences are provided in the Supporting Information). We find that, like the original Mpipi model, the Mpipi-Recharged model achieves excellent agreement with both the experimental *R*_*g*_ (Fig. S2), and phase-diagrams^9,24^ (Fig. 2a–b).

**FIG. 2.**
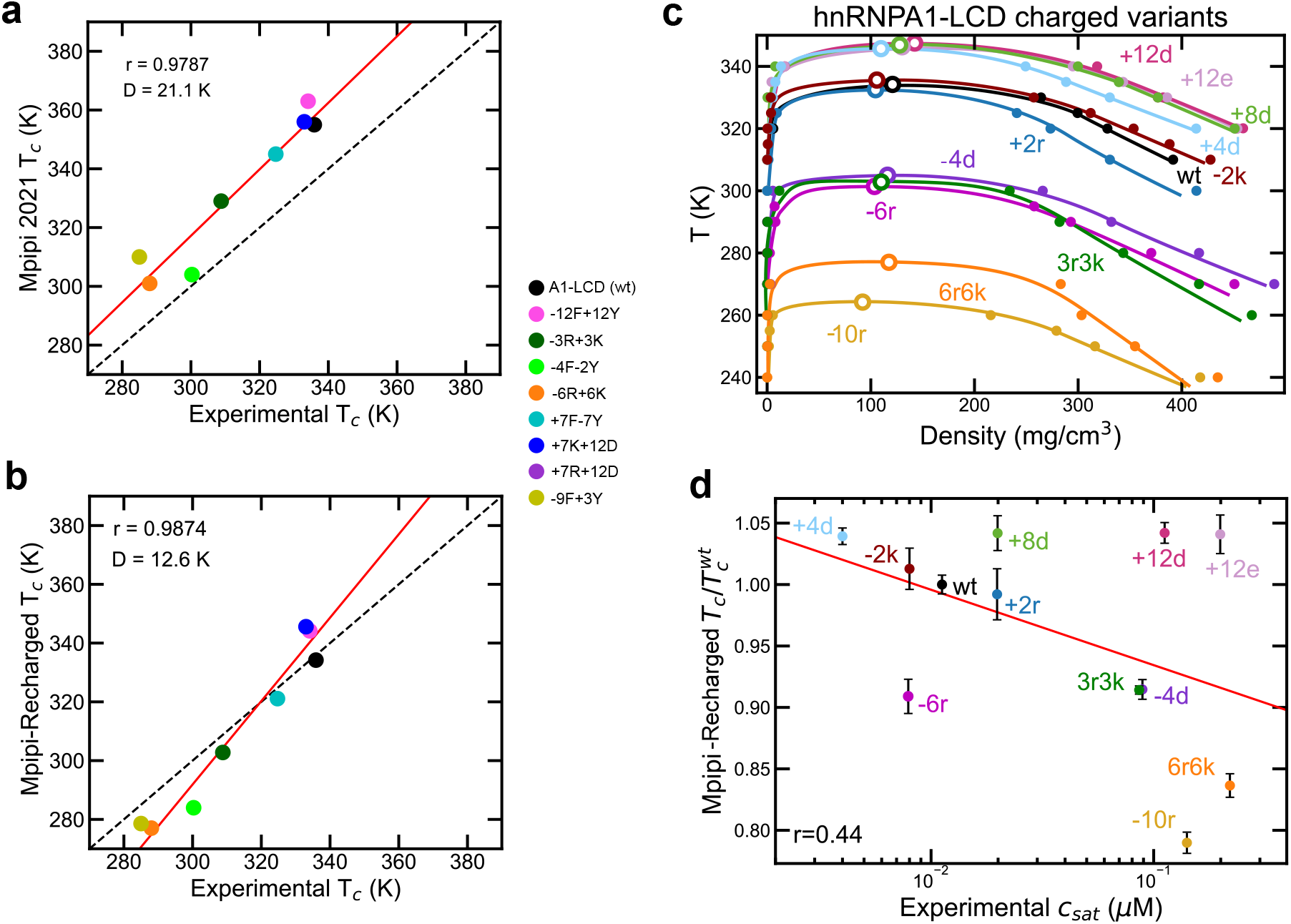
Predicted phase separation behaviour of hnRNPA1-LCD charged variants by the Mpipi-Recharged model. Predicted critical temperature T_*c*_ by Mpipi **a** and Mpipi-Recharged **b** models against experimentally reported critical temperatures for hnRNPA1-LCD charged variants from Ref.^9^. The Pearson correlation coefficient (*r*) and the root mean square deviation from the experimental values (*D*) are displayed for each set of modelling data. **c**Temperature-density phase diagram for hnRNPA1-LCD charged variants. Solid symbols represent the obtained coexistence densities from DC simulations, and open symbols the estimated critical temperature through the law of rectilinear diameters and critical exponents^108^. Continuous lines are included as a guide for the eye. **d**. Predicted critical temperatures by the Mpipi-Recharged model against the experimental saturation concentration reported by Bremer *et al*.^9^ for the hnRNPA1-LCD charged variants. The simulated critical temperature has been normalised by 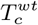 corresponding to wt-hnRNPA1-LCD.

Additionally, we compare critical solution temperatures predicted by the Mpipi-Recharged model (Fig. 2c-d) against the experimental saturation concentrations for various A1-LCD charged variants (*C*_*sat*_)^9^. Such a comparison assumes that the the simulation critical solution temperature and the experimentally measured saturation concentrations are both good proxies of condensate thermodynamic stability^27,78^. Recapitulating the phase behaviour of the A1-LCD charged variants is a stringent test, as mutating just a few charged residues can significantly alter phase behaviour^9^. Charge mutations can either destabilize or promote protein condensation, depending on the characteristics of the introduced residues and the sequence context^9,27,121,122^. Importantly, we find that while the original Mpipi model provides a reasonable correlation between the predicted critical temperatures and the experimental saturation concentrations (Fig. S3), the Mpipi-Recharged model improves the correlation coefficient between these two magnitudes for the charge variants (Fig. 2d).

Capturing the impact of charge blockiness— understood as groups of like-charged residues clustered together along a polymer sequence^123,124^—on the stability of biomolecular condensates with residue-resolution coarse-grained models that consider ions implicitly is a very complex challenge^90^. These models inherently overlook that the distribution of charges within a protein affects the organization and dynamics of ions condensed around the protein, the correlations between these ions, the screening of bare biomolecular charges by these ions, and the entropic gain upon ion release during biomolecular interactions. Despite these limitations, models like Mpipi-Recharged can mitigate some of the effects arising from the lack of an explicit representation of ions through appropriate parameterization. Therefore, we now test how well Mpipi-Recharged performs when predicting the phase behaviour of the wildtype (WT) N-terminal domain (NTD) of the DEAD-Box helicase 4 (DDX4) protein, a highly-charged IDR that presents a high degree of charge blockiness^29^. We compare the behaviour of the WT variant, with a charge-scrambled (CS) variant, which has the same total charge but a different distribution of charged residues, and a arginine-to-lysine (RtoK) and phenylalanine-to-alanine (FtoA) variants. The phase behaviour of these four variants have been studied extensively experimentally^88,102,103^ and with theory and simulations^24,29,90,125^.

Experiments have established that at 100 mM of monovalent salt, the relative propensities of the DDX4 variants to phase separate are as follows: WT *>* CS *>* FtoA *>* RtoK^88,90,102,103^. In addition, experimental phase diagrams in the plane of temperature-vs-concentration at various monovalent salt concentrations have been measured for the WT and CS variants^102^. While the original Mpipi model correctly recovered the relative propensities among the four variants^24,90^ (Fig. S4), our upgraded Mpipi-Recharged model improves the quantitative predictions of the critical solution temperatures for WT and CS in comparison to available experimental values (Fig. 3a)^102^. Additionally, Mpipi-Recharged accurately captures the modulation of the decrease in stability of WT and CS condensates with increasing salt concentration (Fig. 3b).

**FIG. 3.**
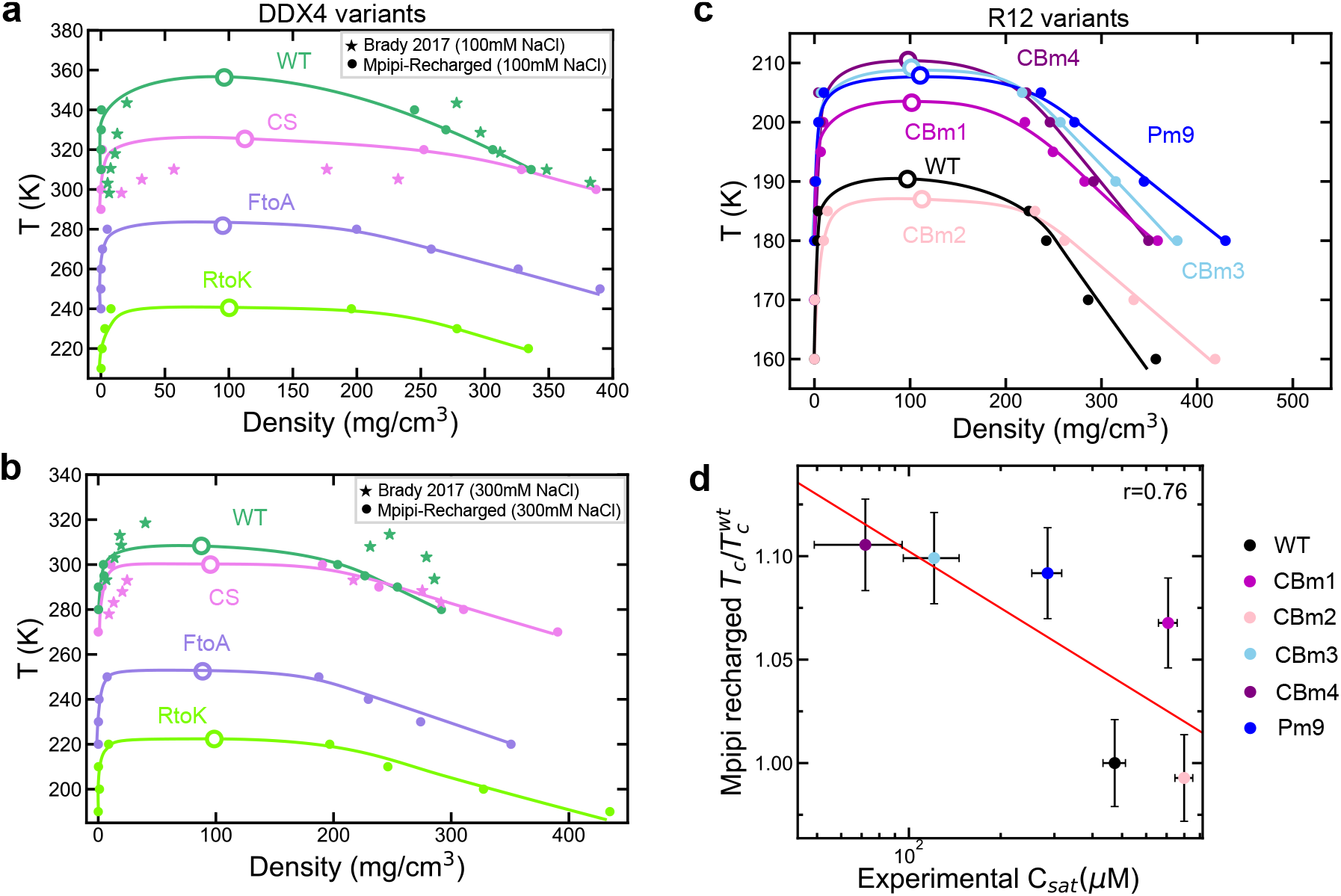
Predicted phase separation behaviour of DDX4 and R12 variants by the Mpipi-Recharged model. Phase diagram in the temperature-density plane (at 100 mM NaCl panel **a** and at 300 mM NaCl panel **b**) for DDX4 variants as studied in Ref.^102^ obtained using DC simulations and the Mpipi-Recharged model. Solid circles indicate the densities of each phase and empty symbols the critical temperature predicted from simulations whereas solid stars represent experimental data from Brady *et al*.^102^ for the WT and CS variants as indicated by the colour code. **c**. Phase diagram in the temperature-density plane for R12 variants with different charged blockiness obtained using DC simulations and the Mpipi-Recharged model. Solid symbols indicate the densities of each phase and empty symbols the critical temperature. **d**. Critical temperature from simulations vs. experimental saturation concentration *C*_*sat*_ for the same R12 variants^46^. The critical temperature is normalised by 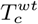 corresponding to the wild-type R12 sequence.

Another phase-separating system where charge block-iness plays a crucial role is the R12 segment of the Ki-67 protein^46^. Experiments analyzing the phase behavior of different R12 variants report a strong correlation between the protein saturation concentration and the degree of charge blockiness^46^). In Fig. 3c, we use Mpipi-Recharged to compute the phase diagram (in the temperature-density plane) for 6 variants of R12 (amino acid sequences provided in the Supporting Information) from Direct Coexistence^84^ simulations. As in the case of DDX4, we find excellent agreement between the predicted propensity to undergo phase separation for each variant (extracted from the T_*c*_ in our simulations) and the experimental saturation concentration reported in Ref.^46^ (Fig. 3d). Overall, these tests confirm the adequacy of coarse-graining charge–charge interactions asymmetrically to improve the description of highly charged condensates.

### C. Mpipi-Recharged captures the regulation of the stability of complex coacervates with salt

Complex coacervates are charge-rich multi-component droplets formed via phase separation of mixtures of oppositely charged polyelectrolyte biomolecules. The stability of complex coacervates is predominantly sustained by heterotypic electrostatic interactions among the different components^7,34–36,53–55^. Not surprisingly, the biophysical properties of complex coacervates, and more generally of charge-rich multi-component condensates, are intricately regulated by changes in their composition and the stoichiometry of the various components. This is because those changes can drastically alter the fine balance between attractive versus repulsive electrostatic interactions ^47,48,51,126–128^. Indeed, monotonic changes in the stoichiometry of the species of multi-component condensates can lead to striking non-linear changes in their stability^26,47,48,51,126–129^. For example, RNA–protein condensates are known to exhibit an RNA-concentration dependent reentrant phase transition, where condensates present low stability at low and high RNA concentrations, and maximum stability at intermediate RNA concentrations^47,48,51,126–128^.

Describing the phase behaviour of complex coacervates is a particularly difficult, yet important, goal for a residue-resolution coarse-grained model. To test the ability of Mpipi-Recharged to probe complex coacervation, we use it to simulate a two-component mixture made of two highly charged proteins: H1 and Prothymosin-*α* (ProT*α*). H1 is a positively charge multi-domain protein, which consists of a short disordered N-terminal tail, a globular domain, and a long (∼100 amino acid) disordered positively charged C-terminal tail.

To treat proteins like H1, we first focus on adapting the Mpipi-Recharged model for multi-domain proteins. Consistent with previous studies, we assume that the secondary structure of globular domains within these multi-domain proteins remains stable throughout the simulations^6,24,63,71,78^. Therefore, we describe globular domains as rigid bodies^6,24,63^. To achieve this, we use an experimental atomistic structure (e.g., from the PDB or AlphaFold) as a reference. The relative positions of all interaction sites within the globular domain are maintained, with beads centered on the *α* carbon^6,24,71,78^.

To account for the burial of amino acids within globular domains, we scale down the interactions of amino acids in these regions^6,24,71,78^. We have done that by titrating the values of two different rescaling factors that reduce the contribution of the Wang–Frenkel potential to the total energy for interactions involving residues within globular domains. The first factor, *δ*_gg_, reduces the Wang–Frenkel interaction strength for globular–globular interactions, while the second factor, *δ*_gIDR_, scales down interactions between a globular domain bead and an IDR bead. During the titration process, we vary the values of *δ*_gg_ and *δ*_gIDR_ from 0.75 to 0.65. We then assess how this titration affects the predicted critical solution temperature estimated by the Mpipi-Recharged model. For the titration, we focus on the phase behaviour of five multi-domain proteins for which experimental saturation concentrations are available; i.e., FUS^51,130,131^, hnRNPA1^132^, HP1^43^, TDP-43^21,134^ (wild-type TDP-43), and a TDP-43 variant with an *α*-helix motif in its C-terminal tail (h-TDP-43)^134^. Detailed information on the atomistic structures used to construct each protein model can be found in the Supporting Material. Additionally, Fig.4a provides a schematic depiction of the model for FUS. Fig. S5 reveals that the best correlation between the experimentally measured saturation concentration and the critical solution temperature predicted with simulations using the Mpipi-Recharged model is reached when *δ*_gg_ = 0.7 and 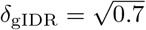 (Fig. 4b).

**FIG. 4.**
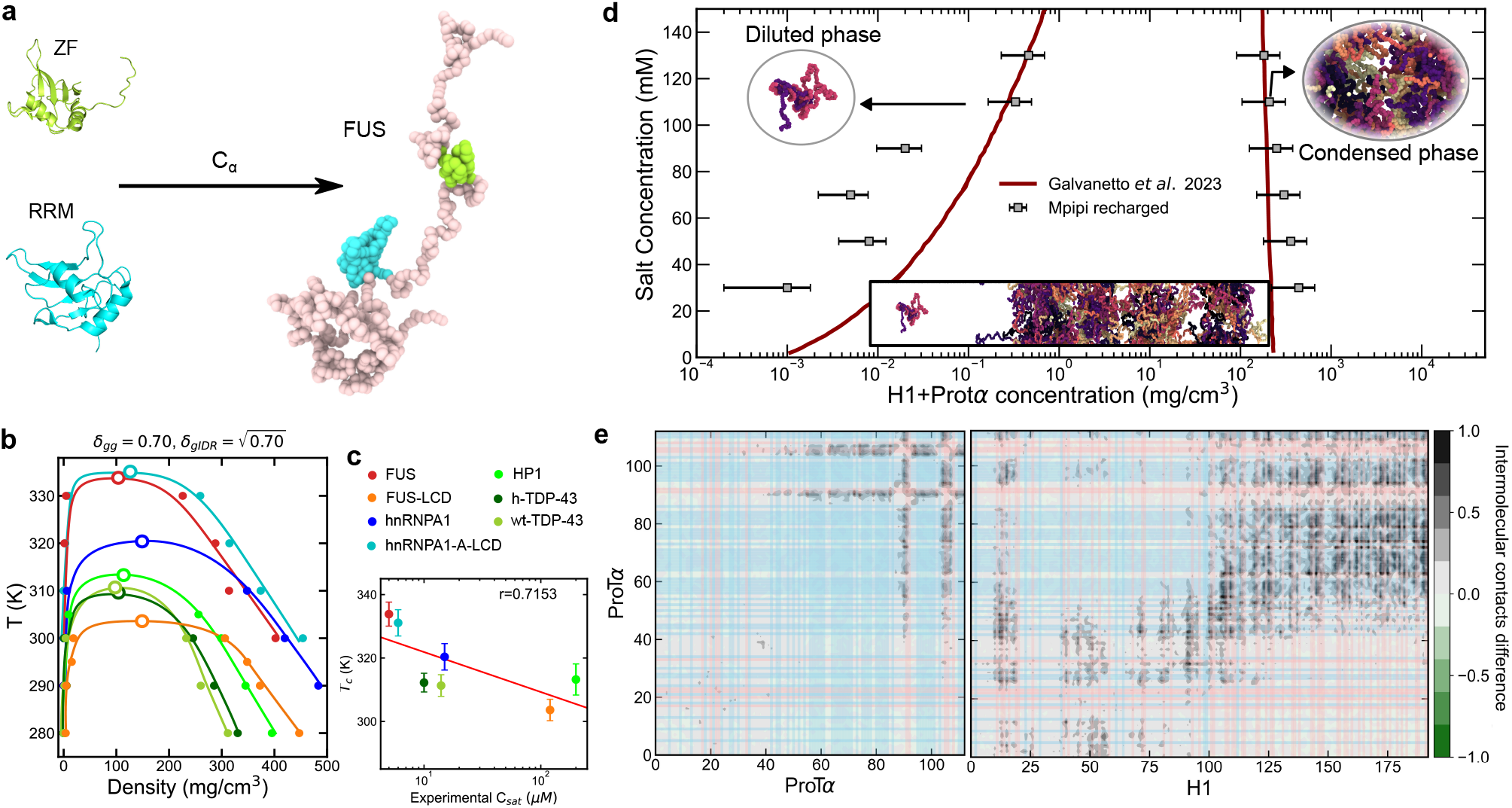
**a**. Schematic representation of the globular domains of FUS from the PDB structures to the coarse-grained representation of the Mpipi-Recharged model. Globular domains (as those depicted by cyan and light green beads for FUS) interactions have been gradually scaled down to optimise the correlation between the model critical temperature and the experimental saturation concentration. **b**. Phase diagram from DC simulations for multi-domain proteins FUS (red), hnRNPA1 (blue), HP1 (lime green), h-TDP-43 (dark green), and wt-TDP-43 (olive green), and intrinsically disordered proteins FUS-LCD (orange), and hnRNPA1-LCD (light blue). Solid symbols indicate the coexistence densities of the two phases and empty symbols display the critical temperature for each system. *δ*_gg_ refers to factor by which Wang–Frenkel interaction strength for globular–globular interactions are rescaled, while *δ*_gIDR_ scales down interactions between a globular domain bead and an IDR bead. **c**. Critical temperature from simulations using the parameters indicated in panel b vs. the experimental saturation concentration *C*_*sat*_ for FUS^51,130,131^, FUS-LCD^131^, hnRNPA1^132^, hnRNPA1-LCD^9,133^, HP1^43^, and TDP-43^21,134^. **d**. Phase diagram of a 1:1.2 H1-ProT*α* mixture (in the salt concentration(KCl)-condensate concentration plane) at *T* = 273*K* predicted by the Mpipi-Recharged model (grey squares). *In vitro* results for the same system reported by Galvanetto *et al*.^7^ are depicted by a red line. Representative snapshots are provided for the simulation slab (bottom) the condensed phase (right) and the dilute phase (left). **e**. Intermolecular contact frequency difference (in number of contacts per residue) between ProT*α*-ProT*α* (Left) and H1-ProT*α* (Right) between the system at 30mM and 130mM of KCl concentration. The thick lines across the panels indicate the positively charged (red) and negatively charged (blue) blocks in ProT*α* and H1 sequences.

After establishing the modelling strategy for multi-domain proteins, we use the Mpipi-Recharged to probe the complex coacervation of the H1:ProT*α* system. Capturing adequately the balance of electrostatic attraction and repulsion in the H1:ProT*α* system is challenging because both proteins have very high net charges: H1 has a net charge of +53e, while ProT*α* has a charge of −44e. Inspired by the experiments of Galvanetto *et al*.^7^, we perform direct coexistence simulations of a H1:ProT*α* mixture with 1:1.2 stoichiometry, which is nearly electroneutral. We vary the monovalent salt concentrations by varying the values of the Yukawa potential screening length (see Methods). From these set of simulations, we compute the phase diagram of the H1:ProT*α* mixture in the plane of monovalent salt concentration versus total biomolecular concentration, which we show in Fig. 4d. Reassuringly, Mpipi-Recharged reproduces the experimental phase diagram (red curve)^7^ with near-quantitative accuracy. Naturally, the simulation predictions of the concentrations of the diluted phase deviate the most from the experimental values because estimating such low concentrations from direct coexistence simulations is inadequate due to the amplification of finite size effects in the diluted phase.

We further investigate the impact of salt concentration on the phase behaviour of the H1:ProT*α* coacervates by evaluating the intermolecular pairwise contact frequencies among the different amino acids in the two proteins. In Fig. 4e we show the variation (in number of contacts per residue; further details provided in Methods) of the number of intermolecular contacts from 130mM to 30mM monovalent salt concentration. Our homotypic intermolecular contact maps for ProT*α*–ProT*α* (Fig. 4e; Left) and heterotypic contact maps for ProT*α*– H1 (Fig. 4e; Right) show a considerable enhancement between oppositely-charged residue contacts (see red (positively charged) and blue (negatively charged) blocks in the chart axes of Figs. 4e and S6) as the monovalent salt concentration decreases. These results reinforce studies showing how the stability of H1:ProT*α* condensates is mainly sustained by electrostatic interactions, and how relatively small variations in salt concentration can strongly regulate their formation and dissolution^6,7,45,68,102,128^.

The excellent agreement between the predictions of the Mpipi-Recharged model with the experimental data for the H1:ProT*α* system further highlights the adequacy of using an asymmetric pair-specific electrostatic potential to improve the description of the phase behaviour of highly charged systems.

### D. Mpipi-Recharged accurately captures the RNA-reentrant behaviour of RNA-binding proteins

Seminal work^47^ by Banerjee and colleagues first discovered that complex coacervates made by short positively-charged synthetic peptides (SR8 and RP3) and single-stranded RNA (polyU) exhibit a RNA-concentration dependent reentrant phase transition. Specifically, they found that the peptide–RNA mixtures only form complex coacervates at intermediate polyU concentrations. When the concentration of polyU was too low (i.e., *C < C*_1_, where *C* is the polyU concentration and *C*_1_ the low concentration threshold) or too high (i.e., *C > C*_2_, where *C*_2_ is the high polyU concentration threshold), phase separation did not take place (Fig. 5a-b; orange diamonds). Furthermore, at intermediate polyU concentrations (*C*_1_ *< C < C*_2_), the stability of the condensates reached a maximum when the stoichiometry of the mixture was near the electroneutral point (*C*_0_). Non-monotonic changes in the stability of multi-component condensates have also been reported for mixtures of FUS-PLD and hnRNAPA1-LCD^26^, and mixtures of PRC1 and RING1B^129^. As in the case of protein–RNA mixtures, in both of these additional examples, associative electrostatic interactions between the two components were identified as the driving forces for the phenomena.

**FIG. 5.**
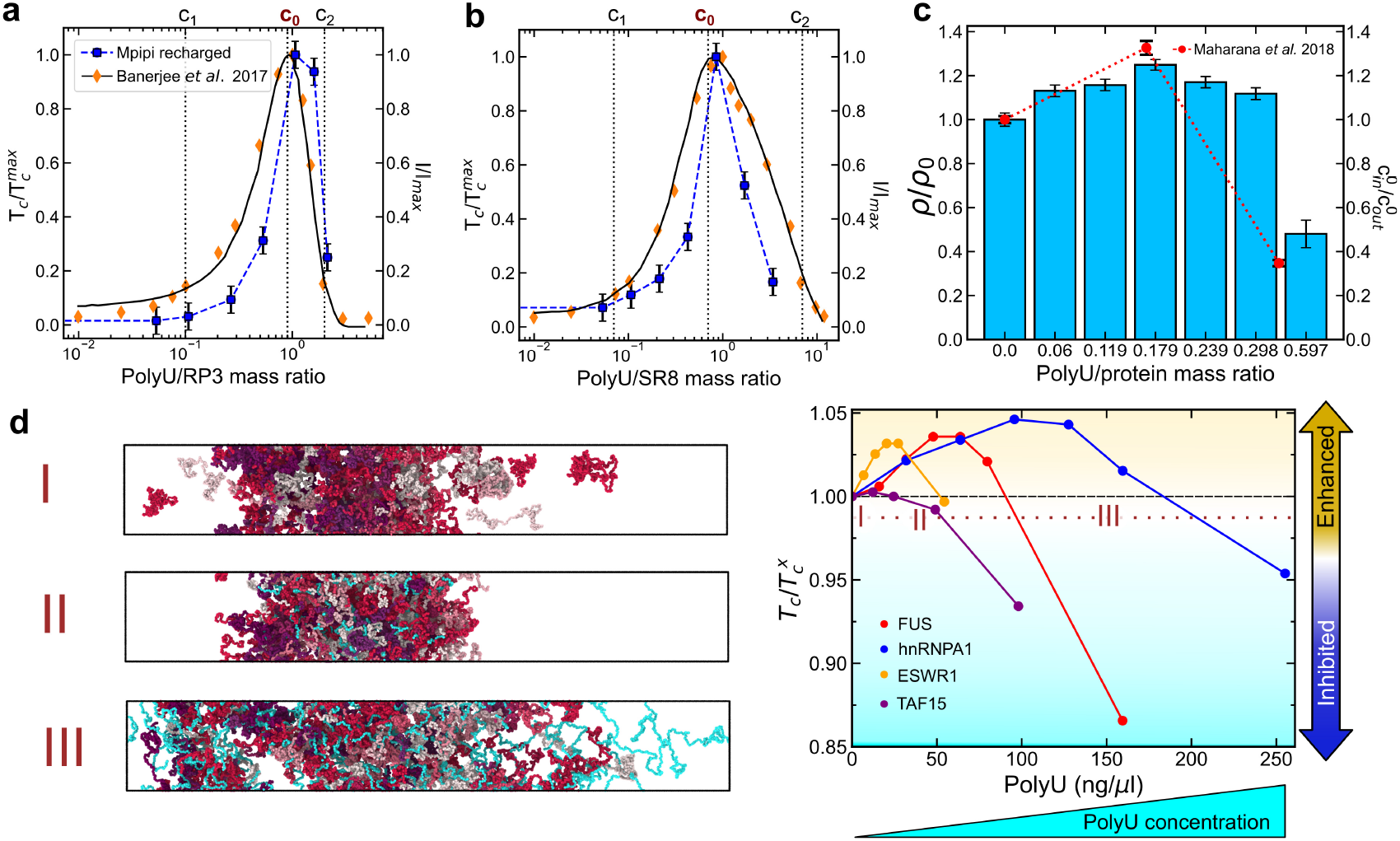
Predictions of the RNA-driven reentrant phase behaviour of protein condensates by the Mpipi-Recharged model. Comparison of simulated critical temperature (blue symbols) with *in vitro* solution turbidity experiments ^47^ (yellow symbols) as a function of the polyU/peptides mass ratio for RP3 (**a**) and SR8 synthetic peptides (**b**). Both simulation critical temperatures (T_*c*_) and fluorescence intensities (I) are normalised by the maximum value of the set. The phase regimes indicated by *C*_1_, *C*_0_ and *C*_2_ are extracted from the work of Banerjee *et al*.^47^ as explained in the text. **c**. Bulk density of polyU/FUS condensates at 300K from NpT simulations (at 0 bar and 300K) as a function of the polyU/protein mass ratio. Red symbols represent the *in vitro* protein partition coefficient for different polyU/FUS mixtures normalised by that of a pure FUS solution as reported by Maharana *et al*.^51^. The computed density from simulations with different concentrations of polyU was renormalized by that of pure FUS condensates at the same conditions. **d**. Right: RNA reentrant behaviour for different multi-domain RBPs (as indicated in the legend) showing the variation in the critical temperature (renormalised by that of pure protein condensates, 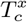) as a function of the polyU concentration. Dashed line sets the separation between promoting and hindering phase separation. Left: Representative snapshots of DC simulations of FUS with different polyU concentrations (as indicated in the left panel), and at a temperature of 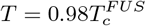 (depicted by the horizontal dotted line in the right panel).

To test whether the Mpipi-Recharged model can reproduce the experimental RNA-driven reentrant phase behaviour of peptide–polyU mixtures observed experimentally by Banerjee et al.^47^, we derive parameters to describe polyU within our Mpipi-Recharged model using a resolution of one-bead per nucleotide. To define the parameters of the Wang–Frenkel and Yukawa potentials for the model, we perform atomistic umbrella sampling simulations of different pairs of amino acids with uracil (U) (Fig. S1). We used the AMBER ff03ws force field both for the amino acids and for the RNA nucleotides^135^. Further details on the simulations and parameter derivations can be found in the Materials and Methods section and the Supporting Information.

Using Mpipi and Mpipi-Recharged, we perform Molecular Dynamics simulations of the condensed liquid phases at constant pressure (0 bar) for SR8–polyU mixtures and RP3–polyU mixtures at various stoichiometries. From these simulations, we estimate the critical solution temperatures (T_*c*_), which provide a quantitative measurement of condensate stability. We have previously demonstrated that pure condensed liquid simulations at constant pressure (e.g., 0 bar) allow for the estimation of critical solution temperatures for multi-component mixtures at fixed stoichiometries, unlike direct coexistence simulations where controlling the condensate composition is difficult^79,85^.

We compare the values of the critical solution temperatures (T_*c*_) predicted by Mpipi (Fig. S7) and Mpipi-Recharged (Fig. 5a-b) against the experimental values of fluorescence intensities reported by Banerjee *et al*.^47^. These values are normalized by the maximum value of T_*c*_, which occurs at concentration *C*_0_. The comparison is made for two different peptide–polyU mixtures as a function of the polyU:peptide ratio, and the results are contrasted with the corresponding experimental values of fluorescence intensities (normalised by maximum intensity at *C*_0_). Figure 5a-b demonstrates that Mpipi-Recharged improves upon the already very good predictions of the original Mpipi model. Notably, Mpipi-Recharged not only recapitulates the RNA-driven reentrant phase behavior of both polyU/RP3 and polyU/SR8—as does the original Mpipi (Figure S7)—but also predicts the concentration thresholds at which maximum stability of the complex coacervates is observed (i.e., *C*_0_) in quantitative agreement with the experimental values. This result underscores the importance of describing attractive and repulsive electrostatic interactions non-symmetrically at the coarse-grained level to achieve better agreement with experimental behaviour.

To further test the performance of Mpipi-Recharged, we examine the behavior of polyU and FUS mixtures, which are also known to exhibit an RNA-driven reentrant phase transition^51,131,136^. Following the experimental setup of Maharana et al.^51^, we perform constant-pressure condensed-liquid simulations of mixtures of 250-nucleotide polyU and the 526-residue FUS protein at various stoichiometries. From these simulations, we measure the densities of the condensed liquids at equilibrium (*ρ*) as a function of the mixture stoichiometry, which directly relates to the stability of the condensates; i.e., higher density implies higher T_*c*_^78,137^, and thus, higher condensate stability. We normalize these condensed liquid densities by dividing them by the density of the pure FUS condensed liquid (*ρ*_0_). The normalized density indicates the relative stability of the different polyU–FUS condensates compared to the pure FUS condensates. We then compare the normalized condensed liquid densities from our simulations with the experimentally measured partition coefficient of FUS^51^. The partition coefficient of FUS is defined as the concentration of FUS inside the condensate (*C*_*in*_) divided by the concentration of FUS in the diluted phase (*C*_*out*_) and, as in the original study, it is presented normalized by the value of the partition coefficient in the pure FUS system 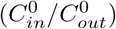. While the original version of Mpipi qualitatively captures the experimental trends^77,85,86^, Mpipi-Recharged, additionally quantitatively predicts the optimal RNA concentration that enhances the stability of polyU/FUS condensates. Moreover, Mpipi-Recharged also correctly predicts the polyU concentration at which the condensates become less stable than those of pure FUS (Fig. 5c).

Finally, we used Mpipi-Recharged to compare the RNA-driven reentrant phase behavior of condensates containing 250-nucleotide polyU with one of the following proteins: EWSR1, TAF15, FUS, or hnRNPA1. For this, we computed the critical temperature of the different protein/polyU condensates as a function of polyU concentration (Fig. 5d). Our simulation results capture the observation of low stability at both low and high polyU concentrations, and high stability at intermediate polyU concentrations for all the four mixtures. Consistently with experimental fluorescent microscopy images, we observe that condensates made from either EWSR1 or TAF15 proteins dissolve at much lower polyU concentrations than hnRNPA1 and FUS condensates. As in the experiments^51^, our simulations show that hnRNPA1 is capable of recruiting much higher amounts of polyU, followed by FUS, before condensate dissolution. The polyU/protein mass ratio threshold at which condensates stability begins to decrease, in all cases, falls very close to the value that produces electroneutral mixtures; that is, ∼11 ng/*μ*l for ESWR1, ∼7 ng/*μ*l for TAF15, ∼25 ng/*μ*l for FUS, and ∼50 ng/*μ*l for hnRNPA1 (for the protein concentration given in ref.^51^). These results suggest that the competition between electrostatic association between proteins and RNA, and the electrostatic repulsion among RNAs as a function of RNA concentration inducing the RNA-driven reentrant phase behaviour observed are well-captured by the Mpipi-Recharged model.

## III. CONCLUSIONS

In this work we introduce the Mpipi-Recharged model, a residue/nucleotide-resolution coarse-grained model for disordered proteins, multi-domain proteins, and mixtures of proteins and single-stranded disordered RNAs. Mpipi-Recharged builds on the success of our original Mpipi model^24^, by improving the description of charge– charge interactions, which are crucial to characterize the behaviour of highly-charged condensates. The Mpipi-Recharged model retains the key advantages of its predecessor: (a) employing the Wang–Frenkel potential^75^ to describe non-electrostatic associative interactions, (b) deriving parameters from a combination of atomistic simulations of amino acid pairs and bioinformatics data, and (c) replacing the commonly used Lorentz-Berthelot combination rules with amino-acid-pair-specific parameters. These features collectively allow the Mpipi models to balance both π-based versus non-π interactions, and arginine versus lysine associative interactions, yielding excellent predictions with experiments^9,24,27^.

A unique feature of the new Mpipi-Recharged model is that it abandons the symmetric description of charge– charge interactions (i.e. the use of a general Debye– Hückel potential for all charged amino acid pairs) in favour of an asymmetric pair-specific Yukawa potential. Such model feature was established based on our atomistic potential of mean force calculations of charged–charged amino acid pairs and charged amino acid/nucleotide pairs, which reveal that at mean-field level, there is a significant asymmetry in the strength of associative versus repulsive electrostatic interactions. Specifically, we observe a substantial reduction (up to −4 kcal/mol) in the free energy when amino acids with opposite charges bind to one another (i.e., R–E, R–D, K–E, and K–D) but negligible changes (less than +1 kcal/mol) when two amino acids of equal charges approach one another (i.e., R–R, K–K, E–E, and D–D). Notably, the decrease in the free energies of arginine upon binding to negatively charged amino acids is much lower than that of lysine, consistent with experimental observations^6,101^ and previous simulations^6,24^.

The striking contrast in the strength of interactions established by opposite-charged versus like-charged species is consistent with differences in the reorientation of amino acid chains as they come closer, changes in ion solvation, and variations in the translational entropy of ions and the solvent. Specifically, as the distance between pairs decreases, the side chains of oppositely charged amino acids, along with the solvating water molecules and surrounding ions, reorganize themselves to maximize (for opposite-charged pairs) Coulomb attraction or minimize (for like-charged pairs) repulsion.

We validate the new Mpipi-Recharge model against experimental single-molecule radii of gyration of approximately 50 IDRs^9,109–120^, achieving near-quantitative agreement in all cases (Fig. S2). Moreover, the model accurately predicts the experimental phase diagrams of hnRNPA1-A-LCD variants *in vitro*^9^ (Fig. 2b), similar to the original Mpipi model^24^.

Addressing the impact of charge blockiness on biomolecular phase behaviour is perhaps one of the most challenging tasks for a coarse-grained model with implicit solvent and ions. These models inherently fail to account for how the distribution of charges within a protein dictates the organization and dynamics of ions surrounding such protein, the correlations between these ions, the screening of biomolecular charges by these ions, and the entropic gains from ion release during biomolecular interactions. Here we show that models like Mpipi-Recharged can alleviate some of these limitations through appropriate parameterization, thus partially compensating for the approximation of physical details, such as the absence of explicit solvent and ion representation. Specifically, we show that the asymmetric coarse-graining of electrostatic forces in Mpipi-Recharged results in excellent agreement with experiments probing the impact of charge blockiness on the stability of DDX4 and the R12 proteins (Fig. 3). An additional stringent test, which we show Mpipi-Recharged excels at, is quantitatively recapitulating the salt-vs-concentration phase diagram of the H1:ProT*α* complex coacervate (Fig. 4d). As in the case of charge blockiness, capturing the complex coacervation of H1:ProT*α* with a coarse-grained model is challenging due to the importance of correctly balancing electrostatic attraction versus repulsion.

Finally, we demonstrate that the Mpipi-Recharged model accurately describes the experimental RNA stoichiometric tresholds at which various protein–RNA condensates exhibit the different regimes of their reentrant phase behaviour^47–49,51^. That is, the model correctly predicts the polyU concentration that maximizes condensate stability for systems including FUS, or short synthetic peptides like RP3 or SR8 with polyU, and the concentration beyond which phase separation is hindered. For RNA-binding proteins such as TAF15, EWSR1, or hnRNPA1, the model matches the range of polyU concentrations that promote and subsequently hinder phase separation^51^ (Fig. 5).

The Mpipi-Recharged model exploits the flexibility of both Wang–Frenkel and Yukawa potentials to include pair-specific parameters to approximate inter-molecular interactions based on the specific chemical makeup of interacting species rather than the absolute charge of each amino acid or the mean hydrophobicity of the pair. This approach enhances the computational tools available for exploring the connection between phase transitions in biomolecular solutions and the physicochemical properties of the constituent biomolecules. Overall, the Mpipi-Recharged model demonstrates how, by compensating for their inherent lack of physical detail with carefully designed energy functions and parameterizations, approximate coarse-grained models can achieve excellent agreement with experimental results. Additionally, it complements experimental studies by offering mechanistic physicochemical insights into biomolecular phase behaviour.

## IV. MATERIALS AND METHODS

### Mpipi-Recharged model for disordered, globular, and multi-domain proteins

Our residue-level coarse-grained model is built based on our original Mpipi model for biomolecular phase separation^138^. Proteins are coarse-grained at amino acid resolution; i.e., each amino acid is represented with a single bead centred on its C_*α*_ atom. The mass of the bead corresponds to the total mass of the amino acid and its molecular diameter (*σ*) is estimated by assuming that the amino acid has a spherical shape with volume equal to its van der Waals volume^139^. Beads that represent amino acids within IDRs/IDPs are connected by stiff harmonic bonds that mantain the mean *C*_*α*_–*C*_*α*_ distance observed experimentally in the backbones of proteins^140^. Globular domains are treated as rigid bodies with the positions of all their amino acid beads being fixed relative to each other using an atomistic structure (e.g. AlphaFold prediction or crystal structure from the Protein Data Bank) as the reference (see Supporting Information for further details).

The total energy of a protein or a protein solution in the Mpipi-Recharged model is the sum of pairwise bonded (*E*_Bonded_) and non-bonded (*E*_Non-bonded_) potentials:

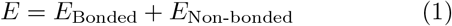

The bonded potential is implemented to mantain amino acids in the protein backbone and is implemented for consecutive amino acids within a protein sequence that belong to the same IDR. However, IDRs are modelled as fully flexible polymers; thus, no energetic penalty for bending or torsion is considered. For bonds between an IDR and a globular domain in a multi-domain protein, we also use the same bonded potential. The bonded pontential is given by:

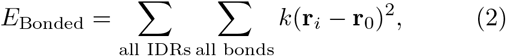

where the equilibrium *C*_*α*_–*C*_*α*_ bond length is *r*_0_ = 3.81Å, as suggested experimentally^140^, and the spring constant, *k* = 9.6 kcal*/*(molÅ^2^), is set sufficiently high to ensure that the equilibrium bond lengths are preserved. The first sum iterates over all IDRs in the system and the second sum iterates over all bonds in each protein, as done before^24^.

The non-bonded potential is calculated for all pair of amino acids *ij*, which are not directly bonded together. This non-bonded potential consist of the sum of a non-electrostatic Wang–Frenkel potential (*E*_WF_) and an electrostatic Yukawa potential (*E*_Y_), as shown in equation (1).

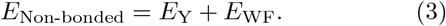

The non-electrostatic Wang–Frenkel potential^75^ (*E*_WF_) is given by

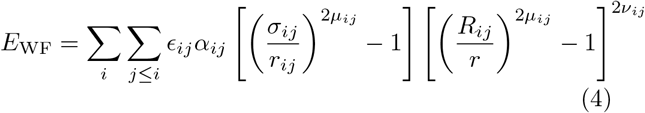

where

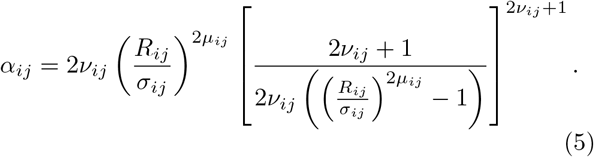

Here *σ*_*ij*_ is the molecular diameter of the pair, defined from the individual molecular diameter of the individual amino acids (*σ*_*i*_ and *σ*_*j*_) using the Lorentz-Berthelot mixing rules: 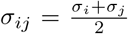). This pair molecular diameter, along with the pair-specific cut-off, *R*_*ij*_ = 3*σ*_*ij*_, define the pairwise distance at which the potential changes from repulsive (*r*_*ij*_ *< σ*_*ij*_) to attractive (*σ*_*ij*_ *> r*_*ij*_ *> R*_*ij*_). As in the original Mpipi model, the depth of the attractive well, −*ϵ*_*ij*_, is set specifically for each pair based on our atomistic potential of mean force calculations and bioinformatics data^88^, rather than defined using Lorentz-Berthelot combination rules. All values of *ϵ*_*ij*_, *σ*_*ij*_, *ν*_*ij*_, and *μ*_*ij*_ are given in Table S4 in the Supporting Information. Abandoning combination rules provides the Mpipi model with unique flexibility in its parameterization, allowing it to correctly balance the much stronger contributions of π-based versus non-π-based associations. For example, unlike what would be possible with Lorentz-Berthelot combination rules, pair-specific parameters can achieve a much larger value of *ϵ*_*ij*_ for both arginine– tyrosine and tyrosine–tyrosine interactions, while keeping that of arginine–arginine sufficiently low to consistently match the atomistic potential of mean force calculations for the three pairs.

The exponents *μ*_*ij*_ and *ν*_*ij*_ are positive integer numbers that modulate the shape of the Wang–Frenkel potential. Specifically, higher values of *μ*_*ij*_ result in a steeper increase in the repulsive part of the potential energy as the distance decreases below *σ*_*ij*_. As in our previous model, we have set *ν*_*ij*_ = 1 for all pairs. However, unlike the original Mpipi model, where *μ*_*ij*_ = 2 for most pairs, we have used higher values of *μ*_*ij*_ ranging from 2 to 12. Consequently, the average value of *μ*_*ij*_ in the Mpipi-Recharged and Mpipi models are ⟨*μ*⟩_*ij*_ = 3.45 and ⟨*μ*⟩_*ij*_ = 2.05, respectively. Using higher values of *μ*_*ij*_ is crucial to avoid interpenetration of beads when values of *ϵ*_*ij*_ are low.

One of the most notable changes in the Mpipi-Recharged model is the description of electrostatic interactions. In the Mpipi-Recharged model, these interactions are described with the Yukawa potential^141^ instead of the Debye–Hückel potential^107^ used originally. The Yukawa potential, given below, is a more general and flexible form of the screened Coulomb potential, given that the strength of interactions can be defined in a pair-specific manner simply by controlling the value of the parameter *A*_*ij*_.

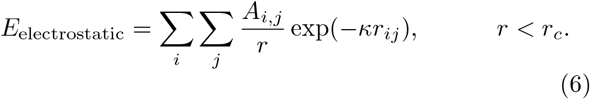

Here *κ* defines the screening length from the concentration of monovalent counterions in solution (*c*_*s*_), according to the expression 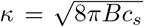, where *B*(*ϵ*_*r*_) = *e*^2^*/*4*πk*_*B*_*Tϵ*_0_*ϵ*_*r*_. The dielectric constant varies with temperature according to the following empiric formula^142^:

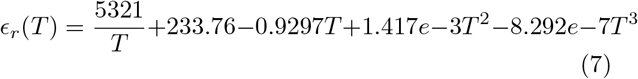

As mentioned earlier, the parameters *A*_*ij*_ of the Yukawa potential ((6)) are determined in a pair-specific manner based on our potential of mean force calculations (Table S3 in the Supporting Information). Our atomistic simulations (Fig. 1) suggest that, at the coarse-grained level, the relative strengths of electrostatic attraction and repulsion among charged amino acid pairs should vary considerably. Therefore, in the Mpipi-Recharged model, we set the Yukawa parameters describing interactions between oppositely charged pairs to be significantly stronger than those among pairs with the same charge. As demonstrated in the results section of this work, this crucial consideration enables the Mpipi-Recharged model to more accurately capture the effects of charge-charge interactions in biomolecular phase separation. Moreover, it outperforms its predecessors for highly-charged systems, while maintaining a low computational cost by avoiding the explicit description of water and ions.

### Model for disordered single-stranded RNA chains

To model disordered single-stranded RNA chains within the Mpipi-Recharged model, we use one bead per nucleotide centered on the phosphorous atom. The total energy for RNA molecules consists on the same bonded and non-bonded terms given in Eq. 1 above. Single-stranded RNA chains are considered as fully flexible polymers, using the stiff harmonic potential *E*_*Bonded*_ given above, with the same spring constant used for proteins but an equilibrium bond length of *r*_0_ = 5.0Å, which is close to the phosphorus–phosphorus distance reported for RNA^143^. As for proteins, non-bonded interactions are defined by the sum of a Wang–Frenkel and Yukawa potentials, with parameters given in Table S3-4 in the Supporting Information.

### Atomistic Umbrella Sampling Molecular Dynamics Simulations to Calculate Potential of Mean Force for Pairs of Amino Acids and Pairs of Amino Acids and Uridine

We perform atomistic umbrella sampling molecular dynamics simulations to estimate the free energy of interactions among charged–charged amino acid pairs and amino acid–Uridine pairs at different values of their centre-of-mass of distances. These simulations were conducted in explicit water and ions, at a salt concentration of 150 mM NaCl, with Na^+^ and Cl^−^ ions. We use the a99sb-disp/PARMBSC1/JC-SPC/E-ion/TIP4P/2005 force field combination^94,95^.

In these simualtions, the heavy atoms are restricted in the direction perpendicular to that of the reaction coordinate using a harmonic potential function with a force constant of 240 kcal/mol/nm^2^. The pairwise center-of-mass distance between the pair of amino acids is varied from around 0.5 nm to 2 nm with 0.05 nm interval, yielding 34 to 40 simulation windows. In each window, we restrain the centre-of-mass distance using a force constant of 1500kcal/mol/nm^2^.

The integration of molecular dynamics is carried out with a 2 fs timestep. The cut-off radius for both dispersive interactions and the real part of electrostatic interactions is 1.4 nm. We use particle mesh Ewald summations to deal with electrostatic interactions. All simulations are run at the constant pressure of p = 1 bar, using an anisotropic Parrinello-Rahman barostat with a relaxation time of 1 ps. To fix the temperature, we employ a velocity-rescale thermostat with a relaxation time of 0.5 ps. Each window is simulated for 10 ns and three independent runs for each system to ensure statistical robustness. To compute potential of mean force calculations, we perform the analysis on the final 9 nanoseconds of each simulation. The final potential of mean force curves are derived using the Weighted Histogram Analysis Method (WHAM)^144^, enhanced with Bayesian bootstrapping. The resulting potential of mean force calculations complement the ones we have estimated previously for the parameterization of our original Mpipi model^24^.

### Calculation of temperature-vs-density phase diagrams via Direct Coexistence simulations

We perform Direct Coexistence simulations^84,145^ to compute the phase diagrams of various protein and protein/RNA solutions in the temperature-vs-concentration plane. These simulations utilize an elongated simulation box to simultaneously simulate the high-density and low-density phases, separated by an interface. The simulation box is a right rectangular prism, with the longer side perpendicular to the interfaces. Periodic boundary conditions are applied across all directions of the simulation box.

Optimising the dimensions of the box is crucial to minimising finite size effects. When defining the simulation box, we adhere to the following guidelines:

1. Use at least 100 proteins per simulation box for single-component systems and 64 for two-component systems.
2. Avoid self-interactions through periodic boundary conditions by ensuring that the short sides of the box are each larger than at least 2*R*_*g*_, where *R*_*g*_ is the radius of gyration of the largest molecule in the mixture within the condensate.
3. Verify that the long dimension of the simulation box is sufficiently long so that the overall density of the biomolecular solution inside the box is approximately 0.1 g/cm^3^. Higher densities can understabilize the condensed phase and may result in a significant underestimation of the critical solution temperature.

Once the simulation setup has been defined, we prepare a preliminary condensed phase at a given temperature by performing an NpT simulation at 1 bar and T=273 K. Then, we place this preliminary condensed phase in the elongated simulation box and run an NVT Direct Co-existence simulation at a given temperature. We use a Langevin thermostat with a relaxation time of 5 ps and a timestep of 10 fs. We perform each simulation for a total of 0.75 to 1.5 *μ*s and verify that equilibrium has been reached by monitoring both the behaviour of the potential energy of the system and the density of the system as as a function of the long side of the simulation box. We then perform a statistical analysis on the last 0.5-1 *μ*s of the simulation post-equilibration. During the analysis, we unwrap the system coordinates on the short sides of the box and re-centre the coordinates of the biomolecules on the centre-of-mass of the condensed phase to calculate the average equilibrium density as a function of the simulation box. If two different densities are detected—i.e., a high density phase and a low density phase—, we average out the densities of the two phases excluding the fluctuations of the interfaces, and plot the value of the two densities (on the x-axis) vs the simulation temperature (y-axis). To construct the coexistence curve of the phase diagram, we vary the temperature and repeat this procedure until only a single homogeneous phase is detected (i.e., when we are above the upper critical solution temperature). For each phase diagram, we estimate the coexistence densities at about 10 different temperatures, reducing the temperature spacing as we get closer to the critical solution temperature (with a maximum grid of 5 K). We then evaluate the critical temperature (*T*_*c*_) and critical density (*ρ*_*c*_) using the law of critical exponents and rectilinear diameters^108^ making sure that for the fit we use only the 3–4 temperature-vs-densities points closer to the critical region that still show sharp interfaces:

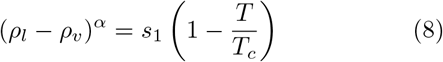

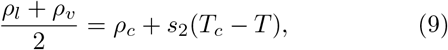

where *ρ*_*l*_ and *ρ*_*v*_ refer to the densities of the condensed and the diluted phases respectively, *s*_1_ and *s*_2_ are fitting parameters, and *α* = 3.06 accounts for the critical exponent of the three dimensional Ising model^108^. All calculations are carried out in LAMMPS^146^. Production runs are typically of the order of 1-2 *μ*s depending on the components of the system and the molecular weight of the protein sequence.

### Estimation of the critical temperature via NpT simulations

To calculate the critical temperature of multi-component mixtures at precise stoichiometries, we run simulations using a cubic homogeneous box of protein/RNA mixtures in the NpT ensemble. This method avoids RNA migration to the diluted phase while ensuring an accurate estimation of the critical temperature^85^. The initial configuration is arranged so that the RNA strands are homogeneously distributed and the density is near bulk conditions *ρ*_*box*_ ∼ 0.3. The temperature and pressure were kept constant using a Nosé–Hoover thermostat^147^ at T=300 K (with 5 ps relaxation time) and a Parrinello–Rahman^148^ isotropic barostat at p=0 bar (with a 5 ps relaxation time), respectively, and a timestep of 10 fs. The critical temperature is calculated running simulations at different temperatures and taking the average of the highest phase separating temperature (with *ρ*_*box*_ ≳ 0.2) and the lowest over-critical temperature (with *ρ*_*box*_ ≲ 0.1). All calculations are carried out in LAMMPS^146^. We note that using this approach, only the coexisting line of the condensed phase can be evaluated, whereas with Direct Coexistence simulations both coexistence lines of the diluted and condensed phase can be measured. However, using the NpT ensemble we ensure precise stoichiometries within condensates for multi-component mixtures.

### Calculations of single-molecule radii of gyration

Single-molecule radii of gyration (*R*_*g*_) are computed for the IDRs listed in the Supporting Information section SI. One replica of each corresponding protein is simulated in a cubic box (with a box size of ∼20 nm) using NVT simulations at the corresponding temperature (see Table S1 in the Supporting Information) for 1.5 *μ*s and a simulation time step of 10 fs. We use a Langevin thermostat with a relaxation time of 5 ps. All calculations are carried out in LAMMPS^146^.

## V. ACKNOWLEDGEMENTS

A. T. is funded by European Research Council (ERC) under the European Union Horizon 2020 research and innovation programme (grant agreement 803326 to R.C.-G.). M.J.M. is funded via the Winton Programme for the Physics of Sustainability. J. R. E acknowledges funding from the Spanish Ministry of Economy and Competitivity (PID2019-105898GA-C22 to J. R. E.) and the Madrid Government (Comunidad de Madrid-Spain) under the Multiannual Agreement with Universidad Politécnica de Madrid in the line Excellence Programme for University Professors, in the context of the V PRICIT (Regional Programme of Research and Technological Innovation). J. R. E. also acknowledges funding from the Roger Ekins Research Fellowship of Emmanuel College, and the Ramon y Cajal fellowship (RYC2021-030937-I to J. R. E.). R.C.-G. acknowledges funding from the ERC under the European Union Horizon 2020 research and innovation programme (grant agreement 803326 R.C.-G.). This project made use of time on high-performance computing granted via the UK High-End Computing Consortium for Biomolecular Simulation, HECBioSim (http://hecbiosim.ac.uk), supported by the Engineering and Physical Sciences Research Council (EPSRC) (grant no. EP/R029407/1 to R.C.-G.). This work also used resources provided by the Cambridge Tier-2 system operated by the University of Cambridge Research Computing Service (http://www.hpc.cam.ac.uk) funded by EP-SRC Tier-2 capital grant EP/P020259/1.

