## Supplementary Material for "Chemically-informed coarse-graining of electrostatic forces in charge-rich biomolecular condensates"

Andrés R. Tejedor

*Yusuf Hamied Department of Chemistry, University of Cambridge, Lensfield Road, Cambridge CB2 1EW, UK and Department of Physical-Chemistry Universidad Complutense de Madrid Av. Complutense s/n, Madrid 28040, Spain*

Anne Aguirre Gonzalez, M. Julia Maristany, Pin Yu Chew, and Kieran Russell

*Yusuf Hamied Department of Chemistry, University of Cambridge, Lensfield Road, Cambridge CB2 1EW, UK*

Jorge Ramirez

*Department of Chemical Engineering, Universidad Politécnica de Madrid, José Gutiérrez Abascal 2, 28006, Madrid, Spain*

Jorge R. Espinosa*

*Department of Physical-Chemistry Universidad Complutense de Madrid Av. Complutense s/n, Madrid 28040, Spain and Maxwell Centre, Cavendish Laboratory, Department of physics, University of Cambridge, JJ Thomson Avenue, Cambridge CB3 0HE, United Kingdom*

Rosana Collepardo-Guevara†

*Yusuf Hamied Department of Chemistry, University of Cambridge, Lensfield Road, Cambridge CB2 1EW, UK and Department of Genetics University of Cambridge Cambridge CB2 3EH, UK*

(Dated: July 26, 2024)

*
†

### SI. SEQUENCES AND PDBS OF THE STUDIED PROTEINS

#### A. Sequences for radii of gyration

Some sequences are such as Prot-$\alpha$ and some hnRNPA1 variants are provided later so we have also calculated their phase behaviour. However, we give all the experimental values of all the radii of gyration studied along with the corresponding reference in Table S1.

**$\alpha$-synuclein:**
MDVFMKGLSK AKEGVVAAAE KTKQGVAEAA GKTKEGVLYV GSKTKEGVVH GVATVAEKTK EQVTNVGGAV
VTGVTAVAQK TVEGAGSIAA ATGFVKKDQL GKNEEGAPQE GILEDMPVDP DNEAYEMPSE EGYQDYEPEA

**ACTR:**
GTQNRPLLRN SLDDLVGPPS NLEGQSDERA LLDQLHTLLS NTDATGLEEI DRALGIPELV NQGQALEPKQ D

**Ash1:**
GASASSSPSP STPTKSGKMR SRSSSPVRPK AYTPSPRSPN YHRFALDSPP QSPRRSSNSS ITKKGSRRSS
GSSPTRHTTR VCV

**FhuA:**
ESAWGPAATI AARQSATGTK TDTPIQKVPQ SISVVTAEEM ALHQPKSVKE ALSYTPGVSV GTRGASNTYD
HLIIRGFAAE GQSQNNYLNG LKLQGNFYND AVIDPYMLER AEIMRGPVSV LYGKSSPGGL LNMVSKRPTT
EPLK

**hNHE1cdt:**
MVPAHKLDSP TMSRARIGSD PLAYEPKEDL PVITIDPASP QSPESVDLVN EELKGKVLGL SRDPAKVAEE
DEDDDGGIMM RSKETSSPGT DDVFTPAPSD SPSSQRIQRC LSDPGPHPEP GEGEPFFPKG Q

**IBB:**
GCTNENANTP AARLHRFKNK GKDSTEMRRR RIEVNVELRK AKKDDQMLKR RNVSSFPDDA TSPLQENRNN
QGTVNWSVDD IVKGINSSNV ENQLQAT

**K10:**
MQTAPVPMPD LKNVKSKIGS TENLKHQPGG GKVQIVYKPV DLSKVTSKCG SLGNIHHKPG GGQVEVKSEK
LDFKDRVQSK IGSLDNITHV PGGGNKKIET HKLTFRENAK AKTDHGAEIV YKSPVVSGDT SPRHLSNVSS
TGSIDMVDSP QLATLADEVS ASLAKQGL

**K16:**
MSSPGSPGTP GSRSRTPSLP TPPTREPKKV AVVRTPPKSP SSAKSRLQTA PVPMPDLKNV KSKIGSTENL
KHQPGGGKVQ IINKKLDLSN VQSKCGSKDN IKHVPGGGSV QIVYKPVDLS KVTSKCGSLG NIHHKPGGGQ
VEVKSEKLDF KDRVQSKIGS LDNITHVPGG GNKKIE

**K17:**
MSSPGSPGTP GSRSRTPSLP TPPTREPKKV AVVRTPPKSP SSAKSRLQTA PVPMPDLKNV KSKIGSTENL
KHQPGGGKVQ IVYKPVDLSK VTSKCGSLGN IHHKPGGGQV EVKSEKLDFK DRVQSKIGSL DNITHVPGGG
NKKIE

**K18:**
MQTAPVPMPD LKNVKSKIGS TENLKHQPGG GKVQIINKKL DLSNVQSKCG SKDNIKHVPG GGSVQIVYKP
VDLSKVTSKC GSLGNIHHKP GGGQVEVKSE KLDFKDRVQS KIGSLDNITH VPGGGNKKIE

**K23:**
MAEPRQEFEV MEDHAGTYGL GDRKDQGGYT MHQDQEGDTD AGLKAEEAGI GDTPSLEDEA AGHVTQARMV
SKSKDGTGSD DKKAKGADGK TKIATPRGAA PPGQKGQANA TRIPAKTPPA PKTPPSSGEP PKSGDRSGYS
SPGSPGTPGS RSRTPSLPTP PTREPKKVAV VRTPPKSPSS AKSRLKKIET HKLTFRENAK AKTDHGAEIV

YKSPVVSGDT SPRHLSNVSS TGSIDMVDSP QLATLADEVS ASLAKQGL

**K25:**
MAEPRQEFEV MEDHAGTYGL GDRKDQGGYT MHQDQEGDTD AGLKAEEAGI GDTPSLEDEA AGHVTQARMV
SKSKDGTGSD DKKAKGADGK TKIATPRGAA PPGQKGQANA TRIPAKTPPA PKTPPSSGEP PKSGDRSGYS
SPGSPGTPGS RSRTPSLPTP PTREPKKVAV VRTPPKSPSS AKSRL

**K27:**
MSSPGSPGTP GSRSRTPSLP TPPTREPKKV AVVRTPPKSP SSAKSRLQTA PVPMPDLKNV KSKIGSTENL
KHQPGGGKVQ IVYKPVDLSK VTSKCGSLGN IHHKPGGGQV EVKSEKLDFK DRVQSKIGSL DNITHVPGGG
NKKIETHKLT FRENAKAKTD HGAEIVY

**K32:**
MSSPGSPGTP GSRSRTPSLP TPPTREPKKV AVVRTPPKSP SSAKSRLQTA PVPMPDLKNV KSKIGSTENL
KHQPGGGKVQ IINKKLDLSN VQSKCGSKDN IKHVPGGGSV QIVYKPVDLS KVTSKCGSLG NIHHKPGGGQ
VEVKSEKLDF KDRVQSKIGS LDNITHVPGG GNKKIETHKL TFRENAKAKT DHGAEIVY

**K44:**
MAEPRQEFEV MEDHAGTYGL GDRKDQGGYT MHQDQEGDTD AGLKAEEAGI GDTPSLEDEA AGHVTQARMV
SKSKDGTGSD DKKAKGADGK TKIATPRGAA PPGQKGQANA TRIPAKTPPA PKTPPSSGEP PKSGDRSGYS
SPGSPGTPGS RSRTPSLPTP PTREPKKVAV VRTPPKSPSS AKSRLQTAPV PMPDLKNVKS KIGSTENLKH
QPGGGKVQIV YKPVDLSKVT SKCGSLGNIH HKPGGGQVEV KSEKLDFKDR VQSKIGSLDN ITHVPGGGNK
KIE

**N49:**
GCQTSRGLFG NNNTNNINNS SSGMNNASAG LFGSKP

**N98:**
GCFNKSFGTP FGGGTGGFGT TSTFGQNTGF GTTSGGAFGT SAFGSSNNTG GLFGNSQTKP GGLFGTSSFS
QPATSTSTGF GFGTSTGTAN TLFGTASTGT SLFSSQNNAF AQNKPTGFGN FGTSTSSGGL FGTTNTTSNP
FGSTSGSLFG P

**NLS:**
ACETNKRKRE QISTDNEAKM QIQEEKSPKK KRKKRSSKAN KPPE

**NSP:**
GCNFNTPQQN KTPFSFGTAN NNSNTTNQNS STGAGAFGTG QSTFGFNNSA PNNTNNANSS ITPAFGSNNT
GNTAFGNSNP TSNVFGSNNS TTNTFGSNSA GTSLFGSSSA QQTKSNGTAG GNTFGSSSLF NNSTNSNTTK
PAFGGLNFGG GNNTTPSSTG NANTSNNLFG ATANAN

**NUL:**
GCGFKGFDTS SSSSNSAASS SFKFGVSSSS SGPSQTLTST GNFKFGDQGG FKIGVSSDSG SINPMSEGFK
FSKPIGDFKF GVSSESKPEE VKKDSKNDNF KFGLSSGLSN PV

**NUS:**
GCPSASPAFG ANQTPTFGQS QGASQPNPPG FGSISSSTAL FPTGSQPAPP TFGTVSSSSQ PPVFGQQPSQ
SAFGSGTTPN

**P53:**
MEEPQSDPSV EPPLSQETFS DLWKLLPENN VLSPLPSQAM DDLMLSPDDI EQWFTEDPGP DEAPRMPEAA
PPVAPAPAAP TPAAPAPAPS WPL

**RNaseA:**
VLLPLLVLVL LLVRVEPSLG KETAAAKFER QHIDSNPSSV SSSNYCNQMM KSRNLTQGRC KPVNTFVHES
LADVQAVCSQ KNVACKNGQT NCYQSYSTMS ITDCRETGSS KYPNCAYKTT QAKKHIIVAC EGNPYVPVHY
DASV

**Sic1:**
GSMTPSTPPR SRGTRYLAQP SGNTSSSALM QGQKTPQKPS QNLVPVTPST TKSFKNAPLL APPNSNMGMT
SPFNGLTSPQ RSPFPKSSVK RT

**SH4-UD:**
MGSNKSKPKD ASQRRRSLEP AENVHGAGGG AFPASQTPSK PASADGHRGP SAAFAPAAAE PKLFGGFNSS DTVTSPQRAG PLAGG

**A1:**
GSMASASSSQ RGRSGSGNFG GGRGGGFGGN DNFGRGGNFS GRGGFGGSRG GGGYGGSGDG YNGFGNDGSN FGGGGSYNDF GNYNNQSSNF GPMKGGNFGG RSSGGSGGGG QYFAKPRNQG GYGGSSSSSS YGSGRRF

**A1-NLS:**
GSMASASSSQ RGRSGSGNFG GGRGGGFGGN DNFGRGGNFS GRGGFGGSRG GGGYGGSGDG YNGFGNDGSN FGGGGSYNDF GNYNNQSSNF GPMKGGNFGG RSSGPYGGGG QYFAKPRNQG GYGGSSSSSS YGSGRRF

**-8F+4Y:**
GSMASASSSQ RGRSGSGNFG GGRGGGYGGN DNGGRGGNYS GRGGFGGSRG GGGYGGSGDG YNGGGNDGSN YGGGGSYNDS GNYNNQSSNF GPMKGGNYGG RSSGGSGGGG QYGAKPRNQG GYGGSSSSSS YGSGRRF

**-9F+6Y:**
GSMASASSSQ RGRSGSGNFG GGRGGGYGGN DNYGRGGNYS GRGGFGGSRG GGGYGGSGDG YNGGGNDGSN YGGGGSYNDS GNYNNQSSNF GPMKGGNYGG RSSGGSGGGG QYGAKPRNQG GYGGSSSSSS YGSGRRY

**+2R:**
GSMASASSSQ RGRSGSGNFG GGRGGGFGGN DNFGRGGNFS GRGGFGGSRG GGGYGGSGDG YNGFRNDGSN FGGGGRYNDF GNYNNQSSNF GPMKGGNFGG RSSGPYGGGG QYFAKPRNQG GYGGSSSSSS YGSGRRF

**+7R:**
GSMASASSSQ RGRSGRGNFG GGRGGGFGGN DNFGRGGNFS GRGGFGGSRG GGRYGGSGDR YNGFGNDGRN FGGGGSYNDF GNYNNQSSNF GPMKGGNFRG RSSGPYGRGG QYFAKPRNQG GYGGSSSSRS YGSGRRF

**-10R+10K:**
GSMASASSSQ KGKSGSGNFG GGKGGGFGGN DNFGKGGNFS GKGGFGGSKG GGGYGGSGDG YNGFGNDGSN FGGGGSYNDF GNYNNQSSNF GPMKGGNFGG KSSGGSGGGG QYFAKPKNQG GYGGSSSSSS YGSGKKF

#### B. hnRNPA1 variants

Below, we give the sequence of hnRNPA1-A-LCD (wild-type) and the different variants we employed.

**wt:**
MASASSSQRG RSGSGNFGGG RGGGFGGNDN FGRGGNFSGR GGFGGSRGGG GYGGSGDGYN GFGNDGSNFG GGGSYNDFGN YNNQSSNFGP MKGGNFGGRS SGPYGGGGQY FAKPRNQGGY GGSSSSSSYG SGRRF

**-3R+3K:**
MASASSSQRG KSGSGNFGGG RGGGFGGNDN FGRGGNFSGR GGFGGSKGGG GYGGSGDGYN GFGNDGSNFG GGGSYNDFGN YNNQSSNFGP MKGGNFGGRS SGGSGGGGQY FAKPRNQGGY GGSSSSSSYG SGRKF

**-4F-2Y:**
MASASSSQRG RSGSGNSGGG RGGGFGGNDN FGRGGNSSGR GGFGGSRGGG GYGGSGDGYN GFGNDGSNSG GGGSSNDFGN YNNQSSNFGP MKGGNFGGRS SGGSGGGGQY SAKPRNQGGY GGSSSSSSSG SGRRF

**-6R+6K:**
MASASSSQKG KSGSGNFGGG RGGGFGGNDN FGKGGNFSGR GGFGGSKGGG GYGGSGDGYN GFGNDGSNFG GGGSYNDFGN YNNQSSNFGP MKGGNFGGKS SGGSGGGGQY FAKPRNQGGY GGSSSSSSYG SGRKF

**+7F-7Y:**
MASASSSQRG RSGSGNFGGG RGGGFGGNDN FGRGGNFSGR GGFGGSRGGG GFGGSGDGFN GFGNDGSNFG GGGSFNDFGN FNNQSSNFGP MKGGNFGGRS SGGSGGGGQF FAKPRNQGGF GGSSSSSSFG SGRRF

**+7K+12D**
MASADSSQRD RDDKGNFGDG RGGGFGGNDN FGRGGNFSDR GGFGGSRGDG KYGGDGDKYN GFGNDGKNFG
GGGSYNDFGN YNNQSSNFDP MKGGNFKDRS SGPYDKGGQY FAKPRNQGGY GGSSSSKSYG SDRRF

**+7R+12D**
MASADSSQRD RDDRGNFGDG RGGGFGGNDN FGRGGNFSDR GGFGGSRGDG RYGGDGDRYN GFGNDGRNFG
GGGSYNDFGN YNNQSSNFDP MKGGNFRDRS SGPYDRGGQY FAKPRNQGGY GGSSSSRSYG SDRRF

**-9F+3Y**
MASASSSQRG RSGSGNFGGG RGGGYGGNDN GGRGGNYSGR GGFGGSRGGG GYGGSGDGYN GGGNDGSNYG
GGGSYNDSGN GNNQSSNFGP MKGGNYGGRS SGGSGGGGQY GAKPRNQGGY GGSSSSSSYG SGRRS

**-12F+12Y**
MASASSSQRG RSGSGNYGGG RGGGYGGNDN YGRGGNYSGR GGYGGSRGGG GYGGSGDGYN GYGNDGSNYG
GGGSYNDYGN YNNQSSNYGP MKGGNYGGRS SGGSGGGGQY YAKPRNQGGY GGSSSSSSYG SGRRY

**-12F+12Y**
MASADSSQRD RDDSGNFGDG RGGGFGGNDN FGRGGNFSDR GGFGGSRGDG GYGGDGDGYN GFGNDGSNFG
GGGSYNDFGN YNNQSSNFDP MKGGNFGDRS SGPYDGGGQY FAKPRNQGGY GGSSSSSSYG SDRRF

**+12D**
MASADSSQRD RDDSGNFGDG RGGGFGGNDN FGRGGNFSDR GGFGGSRGDG GYGGDGDGYN GFGNDGSNFG
GGGSYNDFGN YNNQSSNFDP MKGGNFGDRS SGPYDGGGQY FAKPRNQGGY GGSSSSSSYG SDRRF

**+12E**
MASAESSQRE REESGNFGEG RGGGFGGNDN FGRGGNFSER GGFGGSRGEG GYGGEGDGYN GFGNDGSNFG
GGGSYNDFGN YNNQSSNFEP MKGGNFGERS SGPYEGGGQY FAKPRNQGGY GGSSSSSSYG SERRF

**-4D**
MASASSSQRG RSGSGNFGGG RGGGFGGNGN FGRGGNFSGR GGFGGSRGGG GYGGSGGGYN GFGNSGSNFG
GGGSYNGFGN YNNQSSNFGP MKGGNFGGRS SGPYGGGGQY FAKPRNQGGY GGSSSSSSYG SGRRF

**+4D**
MASASSSQRD RSGSGNFGGG RGGGFGGNDN FGRGGNFSGR GDFGGSRGGG GYGGSGDGYN GFGNDGSNFG
GGGSYNDFGN YNNQSSNFGP MKGGNFGGRS SDPYGGGGQY FAKPRNQGGY GGSSSSSSYD SGRRF

**+8D**
MASASSSQRD RSGSGNFGGG RDGGFGGNDN FGRGDNFSGR GDFGGSRDGG GYGGSGDGYN GFGNDGSNFG
GGGSYNDFGN YNNQSSNFGP MKGGNFGGRS SDPYGGGGQY FAKPRNQDGY GGSSSSSSYD SGRRF

**-10R**
MASASSSQGG SSGSGNFGGG GGGGFGGNDN FGGGGNFSGS GGFGGSGGGG GYGGSGDGYN GFGNDGSNFG
GGGSYNDFGN YNNQSSNFGP MKGGNFGGSS SGPYGGGGQY FAKPGNQGGY GGSSSSSSYG SGGGF

**-6R**
MASASSSQGG RSGSGNFGGG RGGGFGGNDN FGGGGNFSGS GGFGGSRGGG GYGGSGDGYN GFGNDGSNFG
GGGSYNDFGN YNNQSSNFGP MKGGNFGGSS SGPYGGGGQY FAKPGNQGGY GGSSSSSSYG SGGRF

#### C. Multi-domain proteins

Here, we provide the sequence and PDBs corresponding to the analysis of multi-domain proteins. We also include the sequence of FUS-LCD.

**FUS:**
MASNDYTQQA TQSYGAYPTQ PGQGYSQQSS QPYGQQSYSG YSQSTDTSGY GQSSYSSYGQ SQNTGYGTQS
TPQGYGSTGG YGSSQSSQSS YGQQSSYPGY GQQPAPSSTS GSYGSSSQSS SYGQPQSGSY SQQPSYGGQQ
QSYGQQQSYN PPQGYGQQNQ YNSSSGGGGG GGGGGNYGQD QSSMSSGGGS GGGYGNQDQS GGGGSGGYGQ
QDRGGRGRGG SGGGGGGGGG GYNRSSGGYE PRGRGGGRGG RGGMGGSDRG GFNKFGGPRD QGSRHDSEQD
NSDNNTIFVQ GLGENVTIES VADYFKQIGI IKTNKKTGQP MINLYTDRET GKLKGEATVS FDDPPSAKAA

IDWFDGKEFS GNPIKVSFAT RRADFNRGGG NGRGGRGRGG PMGRGGYGGG GSGGGGRGGF PSGGGGGGGQ
QRAGDWKCPN PTCENMNFSW RNECNQCKAP KPDGPGGGPG GSHMGGNYGD DRRGGRGGYD RGGYRGRGGD
RGGFRGGRGG GDRGGFGPGK MDSRGEHRQD RRERPY

**FUS-LCD:**
MASNDYTQQA TQSYGAYPTQ PGQGYSQQSS QPYGQQSYSG YSQSTDTSGY GQSSYSSYGQ SQNTGYGTQS
TPQGYGSTGG YGSSQSSQSS YGQQSSYPGY GQQPAPSSTS GSYGSSSQSS SYGQPQSGSY SQQPSYGGQQ
QSYGQQQSYN PPQGYGQQNQ YNS

**hnRNPA1:**
MSKSESPKEP EQLRKLFIGG LSFETTDESL RSHFEQWGTL TDCVVMRDPN TKRSRGFGFV TYATVEEVDA
AMNARPHKVD GRVVEPKRAV SREDSQRPGA HLTVKKIFVG GIKEDTEEHH LRDYFEQYGK IEVIEIMTDR
GSGKKRGFAF VTFDDHDSVD KIVIQKYHTV NGHNCEVRKA LSKQEMASAS SSQRGRSGSG NFGGGRGGGF
GGNDNFGRGG NFSGRGGFGG SRGGGGYGGS GDGYNGFGND GGYGGGGPGY SGGSRGYGSG GQGYGNQGSG
YGGSGSYDSY NNGGGGGFGG GSGSNFGGGG SYNDFGNYNN QSSNFGPMKG GNFGGRSSGP YGGGGQYFAK
PRNQGGYGGS SSSSSYGSGR RF

**TDP-43:**
MSEYIRVTED ENDEPIEIPS EDDGTVLLST VTAQFPGACG LRYRNPVSQC MRGVRLVEGI LHAPDAGWGN
LVYVVNYPKD NKRKMDETDA SSAVKVKRAV QKTSDLIVLG LPWKTTEQDL KEYFSTFGEV LMVQVKKDLK
TGHSKGFGFV RFTEYETQVK VMSQRHMIDG RWCDCKLPNS KQSQDEPLRS RKVFVGRCTE DMTEDELREF
FSQYGDVMDV FIPKPFRAFA FVTFADDQIA QSLCGEDLII KGISVHISNA EPKHNSNRQL ERSGRFGGNP
GGFGNQGGFG NSRGGGAGLG NNQGSNMGGG MNFGAFSINP AMMAAAQAAL QSSWGMMGML ASQQNQSGPS
GNNQNQGNMQ REPNQAFGSG NNSYSGSNSG AAIGWGSASN AGSGSGFNGG FGSSMDSKSS GWGM

**HP1:**
MGKKTKRTAD SSSSEDEEEY VVEKVLDRRV VKGQVEYLLK WKGFSEEHNT WEPEKNLDCP ELISEFMKKY
KKMKEGENNK PREKSESNKR KSNFSNSADD IKSKKKREQS NDIARGFERG LEPEKIIGAT DSCGDLMFLM
KWKDTDEADL VLAKEANVKC PQIVIAFYEE RLTWHAYPED AENKEKETAK S

The following Protein Data Bank (PDB) codes were used to build the globular structured domains of: FUS (residues from 285–371 (PDB code: 2LCW) and from 422–453 (PDB code: 6G99)), wt-TDP-43 (residues 2-38, 40-49 and 51-79 all included in the same PDB, (PDB code: 5MDI) and from residues 193-267 (PDB code: 1WF0)), h-TDP-43 (additionally to the structured domains of wt-TDP-43, this variant has an $\alpha$–helical domain from residues 307-349 (PBD code: 2N2C)), hnRNPA1 (residues from 8-91 and 103-181 in the same PDB (PDB code: 1L3K)), and HP1$\alpha$ dimer (residues 18-75 (PDB code: 3FDT) and the CSD in residues 110-173 (PDB code: 3I3C)).

#### D. R12 variants

Here, we provide the R12 variants studied in this work and taken from [1].

**Wild-type R12:**
KTTKIACKSP QPDPVDTPAS TKQRPKRNLR KADVEEEFLA LRKRTPSAGK AMDTPKPAVS DEKNINTFVE
TPVQKLDLLG NLPGSKRQPQ TPKEKAEALE DLVGFKELFQ TP

**Pm9:**
KTTKIACKSP QPDPVDEPAS TKQRPKRNLR KADVEEEFLA LRKREPEAGK AMDEPKPAVS DEKNINEFVE
EPVQKLDLLG NLPGEKRQPQ EPKEKAEALE DLVGFKELFQ EP

**CBm-1:**
KTTKIACKSP QPDPVDTPEE EKERPKRELR KEDVEEEFLA LRKRTPSAGK AMDTPKPAVS DEKNINTFVE
TPVQKLDLLG NLPGSKRQPQ TPKEKAEALE DLVGFKELFQ TP

**CBm-2:**
KTTKIACKSP QPDPVDTPAS TQQQPQQNLQ QADVEEEFLA LRKRTPSAGK AMDTPKPAVS DEKNINTFVE
TPVQKLDLLG NLPGSKRQPQ TPKEKAEALE DLVGFKELFQ TP

**CBm-3:**
KTTKIACKSP QPDPVDTPAS TKQRPKRNLR KADVEEEFLA LRKRTPSEEK EVDTPKPEVE DEKEIETFVE TPVEKLDLLG NLPGSKRQPQ TPKEKAEALE DLVGFKELFQ TP

**CBm-4:**
KTTKIACKSP QPDPVDTPAS TKQRPKRNLR KADVEEEFLA LRKRTPSAGK AMDTPKPAVS DEKNINTFVE TPVQKLDLLE ELPESKREPE TPKEKEEELE DLVEFKELFE TP

#### E. H1 and ProT$\alpha$

**H1:**
TENSTSAPAA KPKRAKASKK STDHPKYSDM IVAAIQAEKN RAGSSRQSIQ KYIKSHYKVG ENADSQIKLS IKRLVTTGVL KQTKGVGASG SFRLAKSDEP KKSVAFKKTK KEIKKVATPK KASKPKKAAS KAPTKKPKAT PVKKAKKKLA ATPKKAKKPK TVKAKPVKAS KPKKAKPVKP KAKSSAKRAG KKK

**ProT$\alpha$:**
GPMSDAAVDT SSEITTKDLK EKKEVVEEAE NGRDAPANGN AENEENGEQE ADNEVDEEEE EGGEEEEEEE EGDGEEEDGD EDEEAESATG KRAAEDDEDD DVDTKKQKTD EDD

### SII. URIDINE POTENTIAL MEAN FORCE CALCULATIONS.

We carried out potential mean force (PMF) calculations to assess the energetic interplay between Uridine and different amino acids significant for phase separation.

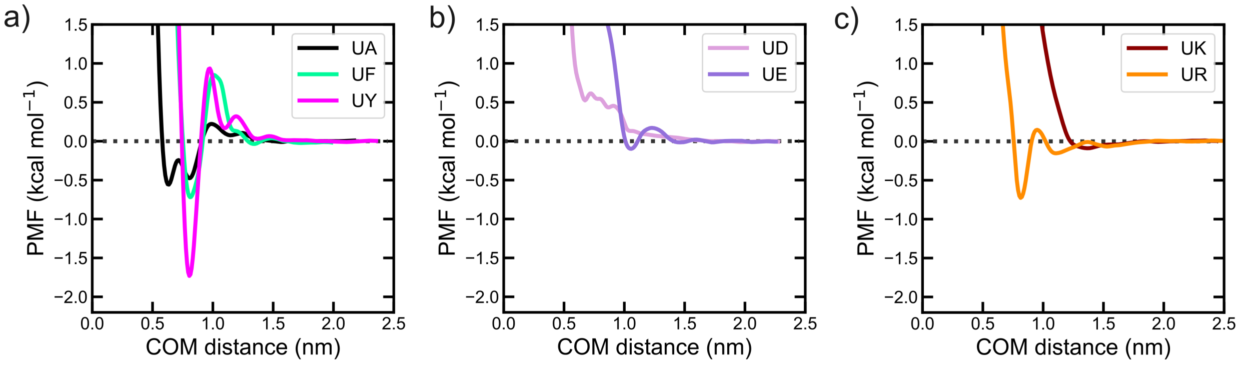


**FIG. S1:** Potential mean force calculations of Uridine nucleotide with different amino-acids including $\pi$ (a), negative (b) and positive (c) residues.

### SIII. SINGLE PROTEIN RADII OF GYRATION.

The radii of gyration provided in Fig. S2 have been calculated as described in the Methods section of the manuscript. The parameter $r$ indicates the linear coefficient parameters and $D$ the deviation.

| | $R_g$ (nm) | $C_{salt}$ (mM) | T (K) | Ref. |
|---|---|---|---|---|
| $\alpha$-synuclein | 3.31 | 185 | 293 | [2] |
| ACTR | 2.51 | 200 | 278 | [3] |
| Ash1 | 2.85 | 150 | 298 | [4] |
| FhuA | 3.34 | 150 | 298 | [5] |
| hNHE1cdt | 3.63 | 200 | 278 | [3] |
| IBB | 3.12 | 162 | 298 | [6] |
| K10 | 4.00 | 150 | 288 | [7] |
| K16 | 3.90 | 150 | 288 | [7] |
| K17 | 3.60 | 150 | 288 | [7] |
| K18 | 3.80 | 150 | 288 | [7] |
| K25 | 4.40 | 150 | 288 | [7] |
| K23 | 4.90 | 150 | 288 | [7] |
| K27 | 3.70 | 150 | 288 | [7] |
| K32 | 4.20 | 150 | 288 | [7] |
| K44 | 5.20 | 150 | 288 | [7] |
| N49 | 1.59 | 162 | 298 | [6] |
| N98 | 2.86 | 162 | 298 | [6] |
| NLS | 2.40 | 162 | 298 | [6] |
| NSP | 4.10 | 162 | 298 | [6] |
| NUL | 3.00 | 162 | 298 | [6] |
| NUS | 2.49 | 162 | 298 | [6] |
| P53 | 2.87 | 108 | 298 | [8] |
| ProT$\alpha$ | 3.79 | 155 | 300 | [9, 10] |
| RNaseA | 3.36 | 150 | 298 | [5] |
| SH4-UD | 2.90 | 217 | 300 | [11] |
| Sic1 | 3.21 | 162 | 298 | [12] |
| A1 | 2.76 | 150 | 298 | [13] |
| A1-NLS | 2.58 | 150 | 298 | [13] |
| -12F+12Y | 2.60 | 150 | 298 | [13] |
| +7F-7Y | 2.72 | 150 | 298 | [13] |
| -9F+6Y | 2.66 | 150 | 298 | [13] |
| -8F+4Y | 2.71 | 150 | 298 | [13] |
| -9F+6Y | 2.68 | 150 | 298 | [13] |
| -10R | 2.67 | 150 | 298 | [13] |
| -6R | 2.57 | 150 | 298 | [13] |
| +2R | 2.62 | 150 | 298 | [13] |
| +7R | 2.71 | 150 | 298 | [13] |
| -3R+3K | 2.63 | 150 | 298 | [13] |
| -6R+6K | 2.79 | 150 | 298 | [13] |
| 10r10k | 2.85 | 150 | 298 | [13] |
| -4D | 2.64 | 150 | 298 | [13] |
| +4D | 2.72 | 150 | 298 | [13] |
| 8D | 2.69 | 150 | 298 | [13] |
| +12D | 2.80 | 150 | 298 | [13] |
| +12E | 2.85 | 150 | 298 | [13] |
| +7K+12D | 2.92 | 150 | 298 | [13] |

**TABLE S1:** Experimental single-protein radii of gyration including the salt concentration ($C_{salt}$), the temperature of the measure, and the reference.

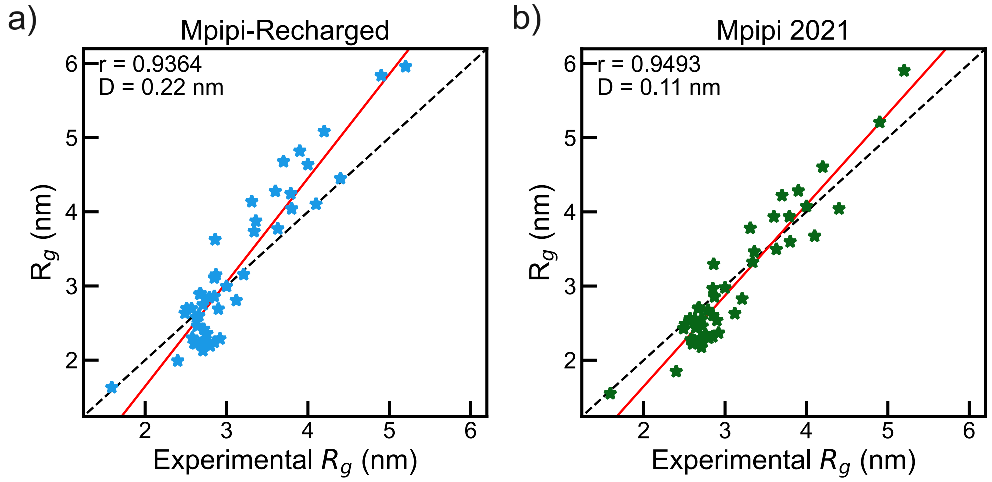


**FIG. S2:** Comparison of the simulated radii of gyration of single-protein using the Mpipi-Recharged (a) and the orginal Mpipi (Mpipi 2021 [14]) (b) with the corresponding experimental value.

### SIV. HNRNPA1 VARIANTS CRITICAL TEMPERATURE

In Table S2 we show the critical temperature obtained from the experiments in [13]. Furthermore, we have calculated the phase diagram of the charged variants of hnRNPA1-LCD and the resulting critical temperature is shown in Fig. S4 for the original Mpipi model [14]. The resulting critical temperature is compared with the experimental saturation concentration obtained in the work of Bremer *et al.* [13].

| Protein | Experimental $T_c$ (K) |
|---|---|
| Wild-type | 336 |
| -12F+12Y | 334 |
| -3R+3K | 309 |
| -4F-2Y | 300 |
| -6R+6K | 288 |
| +7F-7Y | 325 |
| +7K+12D | 333 |
| +7R+12D | 373 |
| -9F+3Y | 285 |

**TABLE S2:** Experimental critical temperature of the different hnRNPA1 variants studied, estimated from the phase diagrams reported by Bremer and co-workers [13].

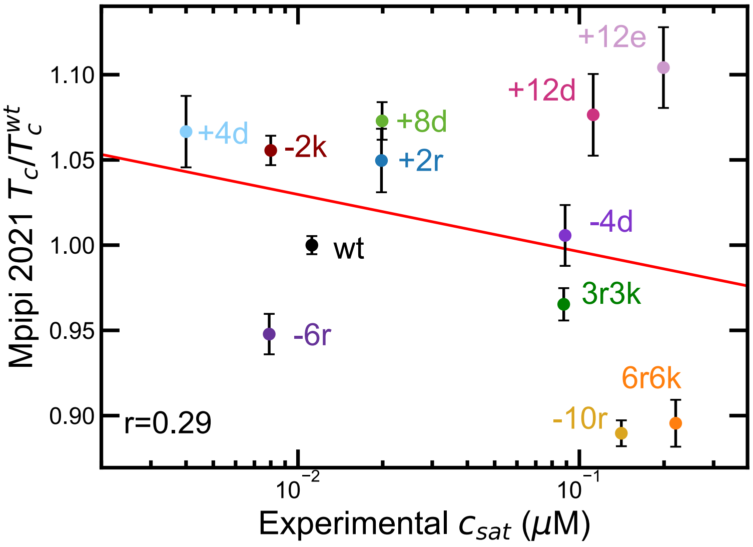


**FIG. S3:** Comparison of the simulated critical temperature using the original Mpipi [14] with the saturation concentration measured in experiments [13].

### SV. DDX4 PHASE DIAGRAMS USING MPIPI MODEL

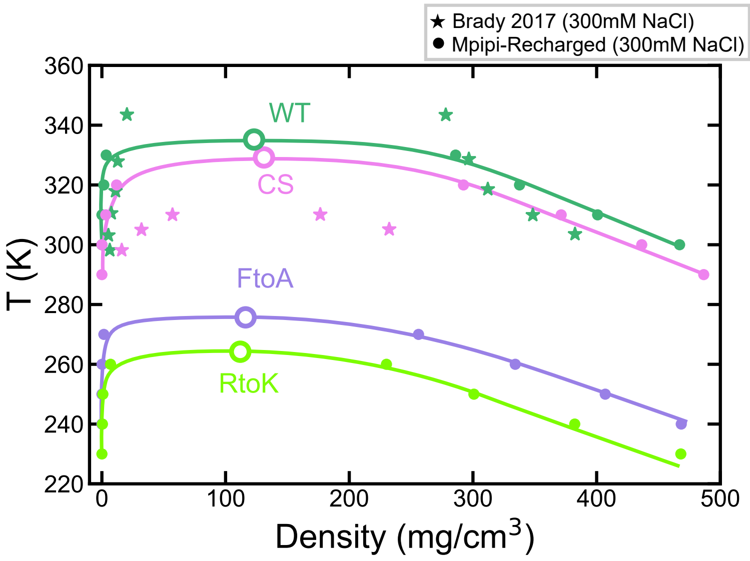


**FIG. S4:** Comparison of the simulated phase diagram using the original Mpipi [14] at [NaCl]=100mM with the experimental phase diagram calculated in the work of Brady *et al.* [15].

### SVI. GLOBULAR PROTEINS ANALYSIS

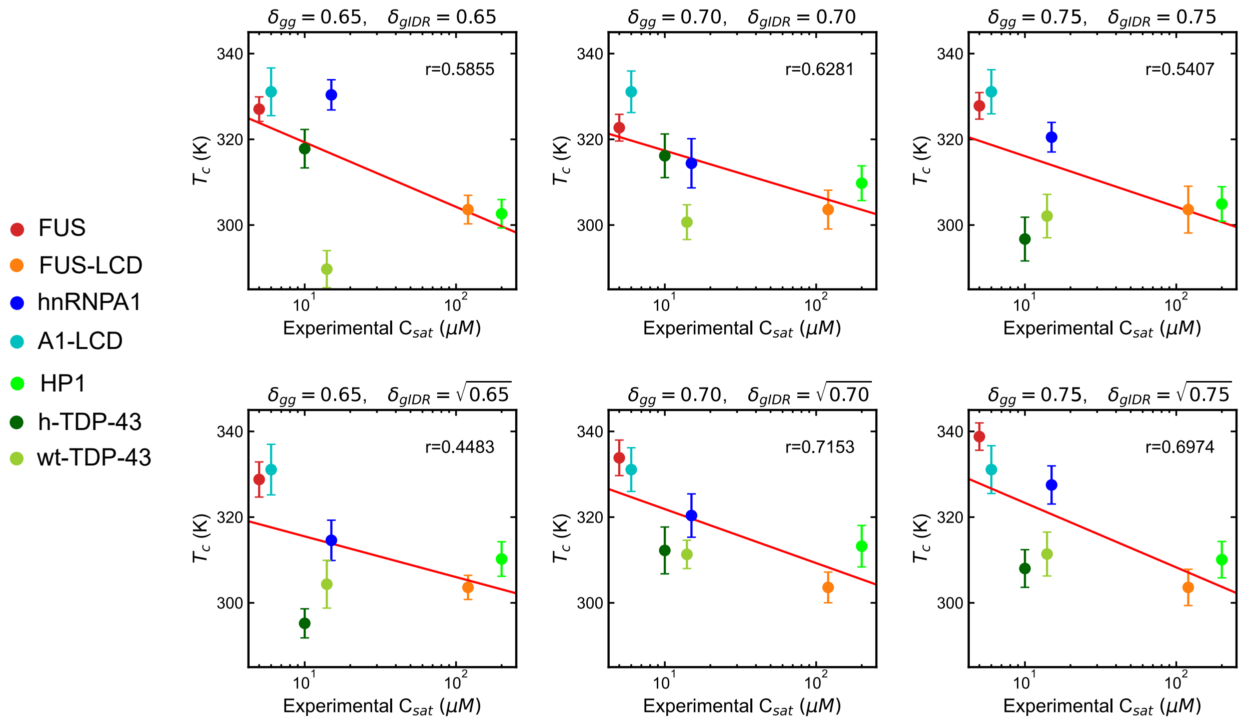


**FIG. S5:** Critical temperature of the studied multi-domain proteins from simulations using different parameters (as indicated in the top of each panel) vs. the experimental saturation concentration $C_{sat}$ for FUS [16–18], FUS-LCD [18], hnRNPA1 [19], hnRNPA1-LCD [13, 20], HP1 [21], and TDP-43 [22, 23].

### SVII. MAPS OF CONTACTS

We calculated the H1-ProT$\alpha$ intermolecular contact maps from DC trajectories at a salt concentration of [NaCl]= 30mM and [NaCl]= 130mM. We used a $\sigma$-dependent cut-off that accounts for the residue specific nature, and it is set to to $1.2\sigma_{ij}$, where $\sigma_{ij}$ accounts for the mean excluded volume of the specific $i^{\text{th}}$ and $j^{\text{th}}$ amino acids. Here we provide all the contact maps between all the components of the H1+ProT$\alpha$ complex coacervate including the position of the charged residues (Fig. S6A-C) and without indicating the charged residues (Fig. S6D-F). The aspect ratio of all the contact maps has been adapted to be a square.

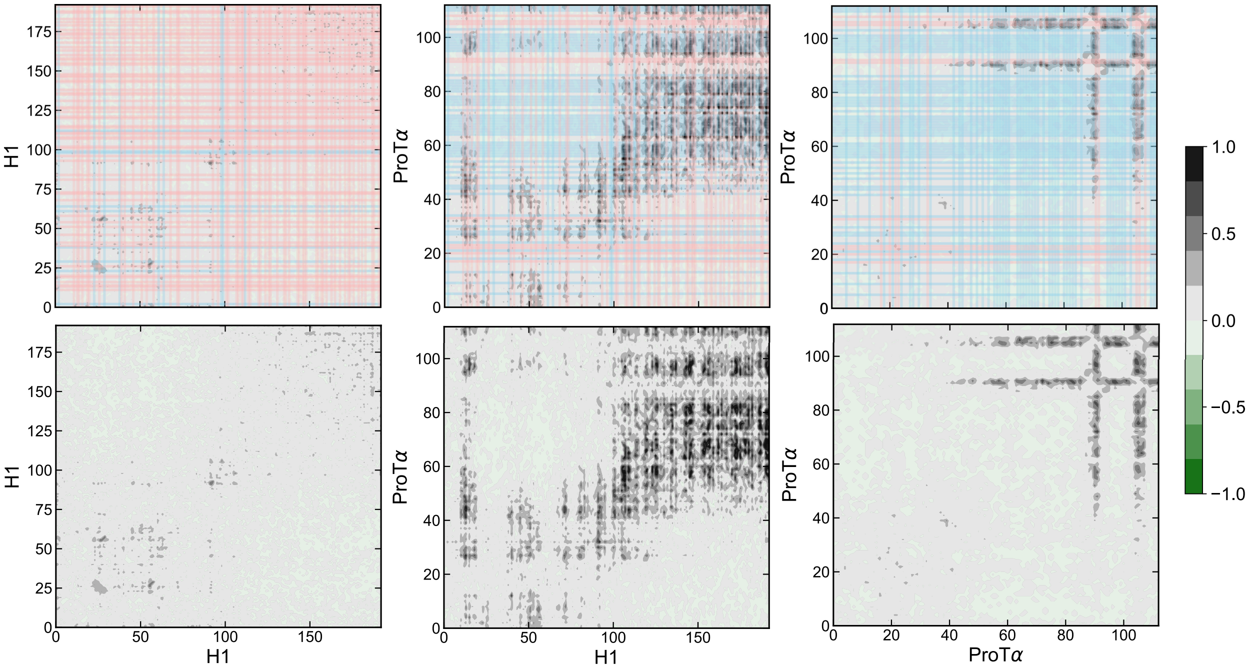


**FIG. S6:** Intermolecular contact frequency difference (in number of contacts per residue) between H1-H1, H1-ProT$\alpha$, and ProT$\alpha$-ProT$\alpha$ (as indicated in the axis labels) between the system at 30mM and 130mM of KCl concentration. Top panels include red lines to indicate the positively charged residues, and blue lines for the negatively charged residues.

### SVIII. RNA RE-ENTRANT BEHAVIOUR USING MPIPI MODEL

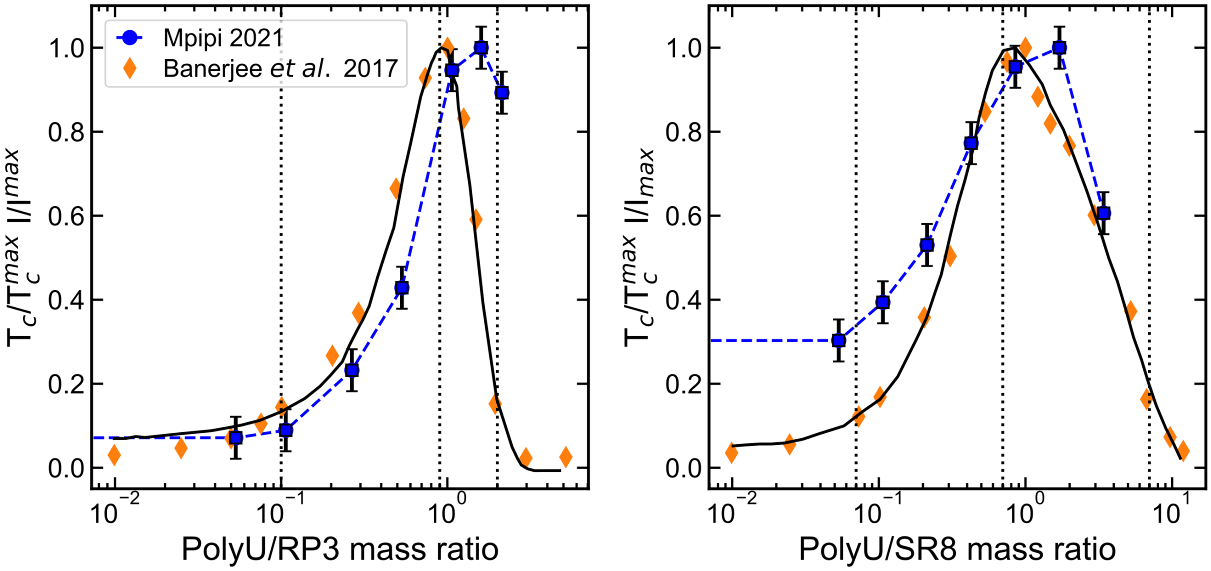


**FIG. S7:** Predictions of the RNA-driven reentrant phase behaviour of protein condensates by the Mpipi model [14]. Comparison of simulated critical temperature (blue symbols) with in vitro solution turbidity experiments [24] (yellow symbols) as a function of the polyU/peptides mass ratio for RP3 (**a**) and SR8 synthetic peptides (**b**). Both simulation critical temperatures ($T_c$) and fluorescence intensities (I) are normalised by the maximum value of the set.

### SIX. MPIPI-RECHARGED PHASE SEPARATION SUMMARY.

We have plot in Fig. S8 a summary figure gathering all the simulated critical temperature vs. the corresponding saturation concentration. The fit to the data gives a significant correlation coefficient of r=0.731 in a wide range of saturation concentrations. The red line represents a linear fit ($y = A \log_{10} x + B$) obtaining the coefficients $A = -53.5K$ and $B = 386K$.

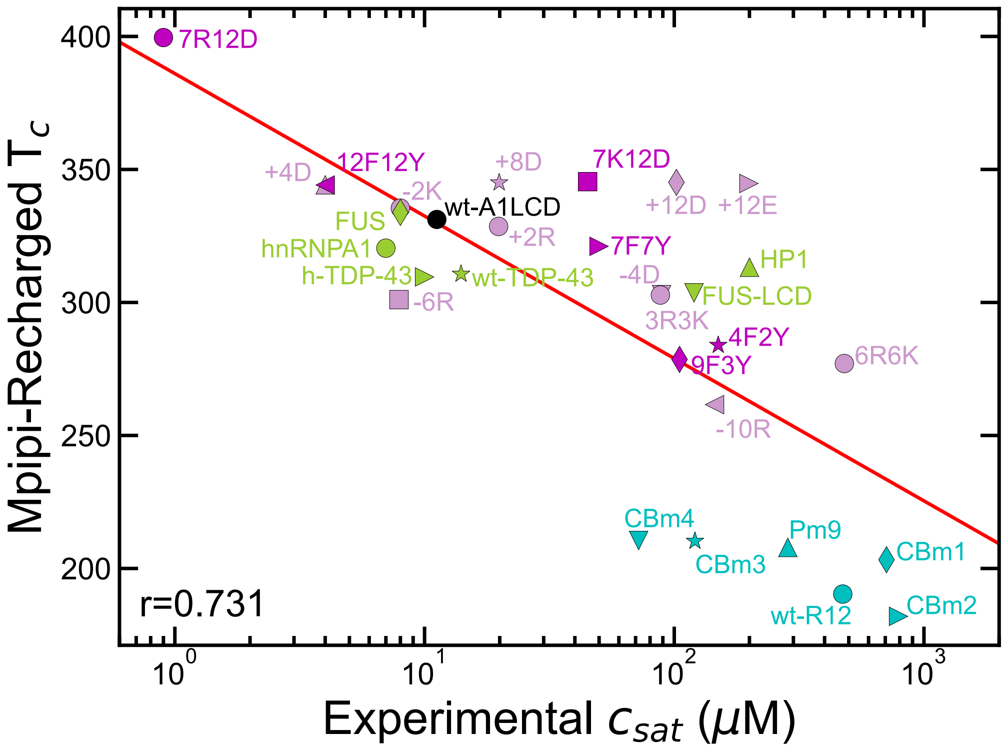


**FIG. S8:** Comparison of our simulated critical temperature with the experimental saturation concentration $C_{sat}$ for all the variants simulated in this work. The colours group the systems in A1-LCD (black), A1-LCD charged variants (pink), other A1-LCD variants (magenta), multi-domain proteins (yellow-green), and R12 variants (cyan). Solid red line provide the fit to the data.

### SX. MPIPI-RECHARGED PARAMETERS

In Table S3 we provide the parameters for the Yukawa interaction for all the charged amino acids. The interaction is given by the parameter $A_{ij}$. The analogous Coulomb potential is given by

$$E_{Coulomb} = \frac{k}{\epsilon}\frac{q_i q_j}{r}, \tag{S1}$$

where $k = 1/4\pi\varepsilon_0$ is the Coulombic constant, $\epsilon$ is the dielectric constant of the medium, and $q_i$, $q_i$ are the charges of the interacting particles. Given this potential, the interaction parameter for the Yukawa potential is calculated as $A_{ij} = k\, q_i q_j/\epsilon$. For the values typically used in previous models, *i.e.* $k = 331.61$ kcal mol$^{-1}$ Å, $\epsilon = 80$, and $q_{i,j} = \pm 1$ (or $q_H = +0.5$ for histidine), the magnitude of the interaction is $|A_{ij}| = 4.145$, (or $|A_{iH}| = 2.073$ when involving histidine). The sign of $A_{ij}$ depends whether the interaction is attractive or repulsive.

| | **R** | **K** | **H** | **D** | **E** |
|---|---|---|---|---|---|
| **R** | +4.00 | +4.00 | +1.47 | -4.91 | -4.93 |
| **K** | +4.00 | +4.00 | +1.48 | -4.33 | -4.34 |
| **H** | +1.47 | +1.48 | +1.04 | -2.46 | -2.47 |
| **D** | -4.91 | -4.33 | -2.46 | +4.00 | +4.00 |
| **E** | -4.93 | -4.34 | -2.47 | +4.00 | +4.00 |

**TABLE S3:** Parameter $A_{ij}$ (in kcal mol$^{-1}$ Å) for the Yukawa potential for each charged pair-residue.

Table of parameters:

| Residue i$^{th}$ | Residue j$^{th}$ | $\epsilon$ (kcal mol$^{-1}$) | $\sigma$ (Å) | $\nu$ | $\mu$ |
|---|---|---|---|---|---|
| **A** | **A** | 0.0912 | 5.2701 | 1 | 4 |
| **A** | **D** | 0.1363 | 5.5468 | 1 | 3 |
| **A** | **E** | 0.1437 | 5.7239 | 1 | 3 |
| **A** | **Y** | 0.3808 | 6.0019 | 1 | 3 |
| **A** | **V** | 0.0529 | 5.7680 | 1 | 4 |
| **A** | **L** | 0.0577 | 5.9021 | 1 | 4 |
| **A** | **Q** | 0.2226 | 5.7740 | 1 | 3 |
| **A** | **W** | 0.5005 | 6.1683 | 1 | 3 |
| **A** | **F** | 0.3582 | 5.9498 | 1 | 3 |
| **A** | **S** | 0.1017 | 5.3414 | 1 | 4 |
| **A** | **H** | 0.3680 | 5.8039 | 1 | 3 |
| **A** | **N** | 0.2169 | 5.5967 | 1 | 3 |
| **A** | **P** | 0.1166 | 5.5381 | 1 | 3 |
| **A** | **C** | 0.1084 | 5.4972 | 1 | 4 |
| **A** | **I** | 0.0484 | 6.0959 | 1 | 4 |
| **C** | **C** | 0.1257 | 5.7244 | 1 | 3 |
| **C** | **I** | 0.0657 | 6.3230 | 1 | 4 |
| **D** | **D** | 0.1379 | 5.8235 | 1 | 3 |
| **D** | **E** | 0.1434 | 6.0006 | 1 | 3 |
| **D** | **Y** | 0.4132 | 6.2786 | 1 | 3 |
| **D** | **V** | 0.0994 | 6.0448 | 1 | 4 |
| **D** | **L** | 0.1040 | 6.1788 | 1 | 4 |
| **D** | **Q** | 0.2632 | 6.0507 | 1 | 3 |
| **D** | **W** | 0.5289 | 6.4450 | 1 | 3 |
| **D** | **F** | 0.3914 | 6.2265 | 1 | 3 |
| **D** | **S** | 0.1465 | 5.6181 | 1 | 3 |
| **D** | **H** | 0.0080 | 6.0807 | 1 | 7 |
| **D** | **N** | 0.2576 | 5.8734 | 1 | 3 |
| **D** | **P** | 0.1608 | 5.8148 | 1 | 3 |
| **D** | **C** | 0.1530 | 5.7739 | 1 | 3 |
| **D** | **I** | 0.0951 | 6.3726 | 1 | 4 |
| **E** | **E** | 0.1489 | 6.1777 | 1 | 3 |
| **E** | **Y** | 0.4202 | 6.4557 | 1 | 3 |
| **E** | **V** | 0.1068 | 6.2218 | 1 | 4 |
| **E** | **L** | 0.1114 | 6.3559 | 1 | 4 |
| **E** | **Q** | 0.2706 | 6.2278 | 1 | 3 |
| **E** | **W** | 0.5360 | 6.6221 | 1 | 3 |
| **E** | **F** | 0.3984 | 6.4036 | 1 | 3 |
| **E** | **S** | 0.1539 | 5.7952 | 1 | 3 |
| **E** | **H** | 0.0080 | 6.2577 | 1 | 7 |
| **E** | **N** | 0.2650 | 6.0505 | 1 | 3 |
| **E** | **P** | 0.1682 | 5.9919 | 1 | 3 |
| **E** | **C** | 0.1604 | 5.9510 | 1 | 3 |
| **E** | **I** | 0.1025 | 6.5497 | 1 | 4 |
| **F** | **F** | 0.6004 | 6.6296 | 1 | 2 |
| **F** | **S** | 0.3681 | 6.0211 | 1 | 3 |
| **F** | **H** | 0.6198 | 6.4837 | 1 | 2 |
| **F** | **N** | 0.4770 | 6.2765 | 1 | 3 |
| **F** | **P** | 0.3822 | 6.2179 | 1 | 3 |
| **F** | **C** | 0.3745 | 6.1770 | 1 | 3 |
| **F** | **I** | 0.3178 | 6.7756 | 1 | 3 |
| **G** | **A** | 0.1321 | 4.9826 | 1 | 3 |
| **G** | **C** | 0.1493 | 5.2097 | 1 | 3 |
| **G** | **D** | 0.1758 | 5.2593 | 1 | 3 |
| **G** | **E** | 0.1832 | 5.4364 | 1 | 3 |
| **G** | **F** | 0.3968 | 5.6623 | 1 | 3 |
| **G** | **G** | 0.1730 | 4.6951 | 1 | 3 |

| | | | | | |
|---|---|---|---|---|---|
| **G** | **H** | 0.4089 | 5.5165 | 1 | 3 |
| **G** | **I** | 0.0893 | 5.8084 | 1 | 4 |
| **G** | **K** | 0.1231 | 5.6832 | 1 | 4 |
| **G** | **L** | 0.0986 | 5.6146 | 1 | 4 |
| **G** | **N** | 0.2578 | 5.3092 | 1 | 3 |
| **G** | **P** | 0.1575 | 5.2506 | 1 | 3 |
| **G** | **Q** | 0.2635 | 5.4865 | 1 | 3 |
| **G** | **R** | 0.3353 | 5.7671 | 1 | 3 |
| **G** | **S** | 0.1426 | 5.0539 | 1 | 3 |
| **G** | **T** | 0.1158 | 5.2921 | 1 | 3 |
| **G** | **V** | 0.0939 | 5.4806 | 1 | 4 |
| **G** | **W** | 0.5393 | 5.8808 | 1 | 3 |
| **G** | **Y** | 0.4195 | 5.7144 | 1 | 3 |
| **H** | **H** | 0.0524 | 6.3378 | 1 | 4 |
| **H** | **N** | 0.4937 | 6.1306 | 1 | 3 |
| **H** | **P** | 0.3934 | 6.0720 | 1 | 3 |
| **H** | **C** | 0.3852 | 6.0311 | 1 | 3 |
| **H** | **I** | 0.3252 | 6.6297 | 1 | 3 |
| **I** | **I** | 0.0057 | 6.9217 | 1 | 12 |
| **K** | **K** | 0.0455 | 6.6713 | 1 | 5 |
| **K** | **T** | 0.0674 | 6.2802 | 1 | 4 |
| **K** | **R** | 0.2167 | 6.7552 | 1 | 3 |
| **K** | **A** | 0.0847 | 5.9707 | 1 | 4 |
| **K** | **D** | 0.0009 | 6.2474 | 1 | 9 |
| **K** | **E** | 0.0009 | 6.4245 | 1 | 9 |
| **K** | **Y** | 0.1842 | 6.7025 | 1 | 3 |
| **K** | **V** | 0.0442 | 6.4687 | 1 | 5 |
| **K** | **L** | 0.0492 | 6.6027 | 1 | 5 |
| **K** | **Q** | 0.2241 | 6.4746 | 1 | 3 |
| **K** | **W** | 0.1984 | 6.8690 | 1 | 3 |
| **K** | **F** | 0.2061 | 6.6504 | 1 | 3 |
| **K** | **S** | 0.0959 | 6.0420 | 1 | 4 |
| **K** | **H** | 0.1663 | 6.5046 | 1 | 3 |
| **K** | **N** | 0.2180 | 6.2973 | 1 | 3 |
| **K** | **P** | 0.1117 | 6.2388 | 1 | 4 |
| **K** | **C** | 0.1030 | 6.1979 | 1 | 4 |
| **K** | **I** | 0.0394 | 6.7965 | 1 | 5 |
| **L** | **L** | 0.0242 | 6.5341 | 1 | 6 |
| **L** | **Q** | 0.1891 | 6.4060 | 1 | 3 |
| **L** | **W** | 0.4687 | 6.8003 | 1 | 3 |
| **L** | **F** | 0.3265 | 6.5818 | 1 | 3 |
| **L** | **S** | 0.0682 | 5.9734 | 1 | 4 |
| **L** | **H** | 0.3345 | 6.4359 | 1 | 3 |
| **L** | **N** | 0.1833 | 6.2287 | 1 | 3 |
| **L** | **P** | 0.0831 | 6.1701 | 1 | 4 |
| **L** | **C** | 0.0749 | 6.1292 | 1 | 4 |
| **L** | **I** | 0.0149 | 6.7279 | 1 | 6 |
| **M** | **A** | 0.0825 | 5.8690 | 1 | 4 |
| **M** | **C** | 0.0998 | 6.0962 | 1 | 4 |
| **M** | **D** | 0.1280 | 6.1457 | 1 | 3 |
| **M** | **E** | 0.1354 | 6.3228 | 1 | 3 |
| **M** | **F** | 0.3507 | 6.5488 | 1 | 3 |
| **M** | **G** | 0.1234 | 5.5815 | 1 | 3 |
| **M** | **H** | 0.3593 | 6.4029 | 1 | 3 |
| **M** | **I** | 0.0398 | 6.6948 | 1 | 5 |
| **M** | **K** | 0.0706 | 6.5696 | 1 | 4 |
| **M** | **L** | 0.0490 | 6.5010 | 1 | 4 |
| **M** | **M** | 0.0739 | 6.4680 | 1 | 4 |
| **M** | **N** | 0.2082 | 6.1957 | 1 | 3 |
| **M** | **P** | 0.1079 | 6.1371 | 1 | 4 |

| | | | | | |
|---|---|---|---|---|---|
| **M** | **Q** | 0.2140 | 6.3729 | 1 | 3 |
| **M** | **R** | 0.2876 | 6.6535 | 1 | 3 |
| **M** | **S** | 0.0931 | 5.9403 | 1 | 4 |
| **M** | **T** | 0.0662 | 6.1785 | 1 | 4 |
| **M** | **V** | 0.0443 | 6.3670 | 1 | 5 |
| **M** | **W** | 0.4923 | 6.7673 | 1 | 3 |
| **M** | **Y** | 0.3727 | 6.6008 | 1 | 3 |
| **N** | **N** | 0.3425 | 5.9234 | 1 | 3 |
| **N** | **P** | 0.2423 | 5.8648 | 1 | 3 |
| **N** | **C** | 0.2341 | 5.8239 | 1 | 3 |
| **N** | **I** | 0.1741 | 6.4225 | 1 | 3 |
| **P** | **P** | 0.1420 | 5.8062 | 1 | 3 |
| **P** | **C** | 0.1338 | 5.7653 | 1 | 3 |
| **P** | **I** | 0.0738 | 6.3639 | 1 | 4 |
| **Q** | **Q** | 0.3540 | 6.2779 | 1 | 3 |
| **Q** | **W** | 0.6252 | 6.6722 | 1 | 2 |
| **Q** | **F** | 0.4824 | 6.4537 | 1 | 3 |
| **Q** | **S** | 0.2332 | 5.8453 | 1 | 3 |
| **Q** | **H** | 0.4994 | 6.3078 | 1 | 3 |
| **Q** | **N** | 0.3483 | 6.1006 | 1 | 3 |
| **Q** | **P** | 0.2480 | 6.0420 | 1 | 3 |
| **Q** | **C** | 0.2399 | 6.0011 | 1 | 3 |
| **Q** | **I** | 0.1799 | 6.5998 | 1 | 3 |
| **R** | **R** | 0.1507 | 6.8391 | 1 | 3 |
| **R** | **A** | 0.2959 | 6.0546 | 1 | 3 |
| **R** | **D** | 0.0067 | 6.3313 | 1 | 7 |
| **R** | **E** | 0.0069 | 6.5084 | 1 | 7 |
| **R** | **Y** | 0.8042 | 6.7863 | 1 | 2 |
| **R** | **V** | 0.2591 | 6.5525 | 1 | 3 |
| **R** | **L** | 0.2636 | 6.6866 | 1 | 3 |
| **R** | **Q** | 0.4225 | 6.5585 | 1 | 3 |
| **R** | **W** | 0.9341 | 6.9528 | 1 | 2 |
| **R** | **F** | 0.7170 | 6.7343 | 1 | 2 |
| **R** | **S** | 0.3061 | 6.1259 | 1 | 3 |
| **R** | **H** | 0.2083 | 6.5884 | 1 | 3 |
| **R** | **N** | 0.4170 | 6.3812 | 1 | 3 |
| **R** | **P** | 0.3204 | 6.3226 | 1 | 3 |
| **R** | **C** | 0.3125 | 6.2817 | 1 | 3 |
| **R** | **I** | 0.2547 | 6.8804 | 1 | 3 |
| **S** | **S** | 0.1123 | 5.4127 | 1 | 4 |
| **S** | **H** | 0.3785 | 5.8752 | 1 | 3 |
| **S** | **N** | 0.2274 | 5.6680 | 1 | 3 |
| **S** | **P** | 0.1271 | 5.6094 | 1 | 3 |
| **S** | **C** | 0.1190 | 5.5685 | 1 | 3 |
| **S** | **I** | 0.0590 | 6.1672 | 1 | 4 |
| **T** | **T** | 0.0586 | 5.8891 | 1 | 4 |
| **T** | **R** | 0.2802 | 6.3641 | 1 | 3 |
| **T** | **A** | 0.0749 | 5.5796 | 1 | 4 |
| **T** | **D** | 0.1206 | 5.8563 | 1 | 3 |
| **T** | **E** | 0.1280 | 6.0334 | 1 | 3 |
| **T** | **Y** | 0.3654 | 6.3113 | 1 | 3 |
| **T** | **V** | 0.0366 | 6.0775 | 1 | 5 |
| **T** | **L** | 0.0414 | 6.2116 | 1 | 5 |
| **T** | **Q** | 0.2063 | 6.0835 | 1 | 3 |
| **T** | **W** | 0.4850 | 6.4778 | 1 | 3 |
| **T** | **F** | 0.3428 | 6.2593 | 1 | 3 |
| **T** | **S** | 0.0854 | 5.6509 | 1 | 4 |
| **T** | **H** | 0.3517 | 6.1134 | 1 | 3 |
| **T** | **N** | 0.2006 | 5.9062 | 1 | 3 |
| **T** | **P** | 0.1003 | 5.8476 | 1 | 4 |

| | | | | | |
|---|---|---|---|---|---|
| **T** | **C** | 0.0921 | 5.8067 | 1 | 4 |
| **T** | **I** | 0.0321 | 6.4054 | 1 | 5 |
| **V** | **V** | 0.0147 | 6.2660 | 1 | 6 |
| **V** | **L** | 0.0194 | 6.4000 | 1 | 6 |
| **V** | **Q** | 0.1844 | 6.2719 | 1 | 3 |
| **V** | **W** | 0.4642 | 6.6663 | 1 | 3 |
| **V** | **F** | 0.3220 | 6.4478 | 1 | 3 |
| **V** | **S** | 0.0635 | 5.8393 | 1 | 4 |
| **V** | **H** | 0.3297 | 6.3019 | 1 | 3 |
| **V** | **N** | 0.1786 | 6.0947 | 1 | 3 |
| **V** | **P** | 0.0784 | 6.0361 | 1 | 4 |
| **V** | **C** | 0.0702 | 5.9952 | 1 | 4 |
| **V** | **I** | 0.0102 | 6.5938 | 1 | 7 |
| **W** | **W** | 0.8031 | 7.0666 | 1 | 2 |
| **W** | **F** | 0.7703 | 6.8481 | 1 | 2 |
| **W** | **S** | 0.5105 | 6.2396 | 1 | 3 |
| **W** | **H** | 0.7632 | 6.7022 | 1 | 2 |
| **W** | **N** | 0.6198 | 6.4950 | 1 | 2 |
| **W** | **P** | 0.5246 | 6.4364 | 1 | 3 |
| **W** | **C** | 0.5169 | 6.3955 | 1 | 3 |
| **W** | **I** | 0.4599 | 6.9941 | 1 | 3 |
| **Y** | **Y** | 0.6458 | 6.7336 | 1 | 2 |
| **Y** | **V** | 0.3447 | 6.4998 | 1 | 3 |
| **Y** | **L** | 0.3492 | 6.6339 | 1 | 3 |
| **Y** | **Q** | 0.5050 | 6.5057 | 1 | 3 |
| **Y** | **W** | 0.7929 | 6.9001 | 1 | 2 |
| **Y** | **F** | 0.6231 | 6.6816 | 1 | 2 |
| **Y** | **S** | 0.3908 | 6.0732 | 1 | 3 |
| **Y** | **H** | 0.6424 | 6.5357 | 1 | 2 |
| **Y** | **N** | 0.4996 | 6.3285 | 1 | 3 |
| **Y** | **P** | 0.4048 | 6.2699 | 1 | 3 |
| **Y** | **C** | 0.3971 | 6.2290 | 1 | 3 |
| **Y** | **I** | 0.3404 | 6.8277 | 1 | 3 |
| **M** | **U** | 0.1723 | 7.3190 | 1 | 3 |
| **G** | **U** | 0.2007 | 6.4326 | 1 | 3 |
| **K** | **U** | 0.0972 | 7.4207 | 1 | 3 |
| **T** | **U** | 0.1679 | 7.0295 | 1 | 3 |
| **R** | **U** | 0.3949 | 7.5045 | 1 | 3 |
| **A** | **U** | 0.1772 | 6.7200 | 1 | 3 |
| **D** | **U** | 0.1920 | 6.9968 | 1 | 3 |
| **E** | **U** | 0.1953 | 7.1738 | 1 | 3 |
| **Y** | **U** | 0.6621 | 7.4518 | 1 | 3 |
| **V** | **U** | 0.1553 | 7.2180 | 1 | 3 |
| **L** | **U** | 0.1580 | 7.3520 | 1 | 3 |
| **Q** | **U** | 0.2527 | 7.2239 | 1 | 3 |
| **W** | **U** | 0.4276 | 7.6183 | 1 | 3 |
| **F** | **U** | 0.3483 | 7.3998 | 1 | 3 |
| **S** | **U** | 0.1833 | 6.7913 | 1 | 3 |
| **H** | **U** | 0.1661 | 7.2539 | 1 | 3 |
| **N** | **U** | 0.2494 | 7.0467 | 1 | 3 |
| **P** | **U** | 0.1918 | 6.9881 | 1 | 3 |
| **C** | **U** | 0.1872 | 6.9472 | 1 | 3 |
| **I** | **U** | 0.1527 | 7.5458 | 1 | 3 |
| **U** | **U** | 0.1100 | 8.1700 | 1 | 3 |

**TABLE S4:** Parameters of the Mpipi-Recharged

[17] J. Wang, J.-M. Choi, A. S. Holehouse, H. O. Lee, X. Zhang, M. Jahnel, S. Maharana,

R. Lemaitre, A. Pozniakovsky, D. Drechsel, *et al.*, "A molecular grammar governing the driving forces for phase separation of prion-like RNA binding proteins," *Cell*, vol. 174, no. 3, pp. 688–699, 2018.
